# Membrane Bound O-Acyltransferase 7 (MBOAT7)-Driven Lysophosphatidylinositol (LPI) Acylation in Adipocytes Contributes to Systemic Glucose Homeostasis

**DOI:** 10.1101/2022.05.19.492632

**Authors:** William Massey, Venkateshwari Varadharajan, Rakhee Banerjee, Amanda L. Brown, Anthony J. Horak, Rachel C. Hohe, E. Ricky Chan, Calvin Pan, Renliang Zhang, Daniela S. Allende, Aldons J. Lusis, J. Mark Brown

**Affiliations:** Department of Cardiovascular and Metabolic Sciences, Lerner Research Institute Cleveland Clinic, Cleveland, OH 44195, USA; Center for Microbiome and Human Health, Lerner Research Institute, Cleveland Clinic, Cleveland, OH 44195, USA; Department of Anatomical Pathology, Cleveland Clinic, Cleveland, OH, USA; Proteomics and Metabolomics Core, Lerner Research Institute, Cleveland Clinic, Cleveland, OH 44195, USA; Institute for Computational Biology, Case Western Reserve University, Cleveland, OH 44106; Departments of Medicine, Microbiology, and Human Genetics, University of California Los Angeles, Los Angeles, CA 90095, USA

**Keywords:** Non-alcoholic fatty liver disease, obesity, metabolism, diabetes

## Abstract

Non-alcoholic fatty liver disease (NAFLD) is becoming increasingly common and is a leading cause of end stage liver diseases such as cirrhosis and hepatocellular carcinoma. The rise in NAFLD closely parallels the global epidemic of obesity and type 2 diabetes mellitus (T2DM), and there is a clear interrelationship between abnormal lipid metabolism, insulin resistance, and NAFLD progression. Several genetic loci have been identified as contributors to NAFLD progression, all of which are consistently linked to abnormal lipid metabolic processes in the liver. The common loss-of-function variant rs641738 (C>T) near the gene encoding Membrane-Bound O-Acyltransferase 7 (*MBOAT7*) is associated with increased susceptibility to NAFLD as well as the entire spectrum of NAFLD progression. The *MBOAT7* gene encodes a lipid metabolic enzyme that is capable of esterifying polyunsaturated fatty acyl-CoAs to LPI substrates to generate phosphatidylinositol (PI) lipids. We previously showed that antisense oligonucleotide (ASO)-mediated knockdown of *Mboat7* in mice promoted high fat diet-induced hepatic steatosis, hyperinsulinemia, and systemic insulin resistance (Helsley et al., 2019). Thereafter, other groups showed that hepatocyte-specific genetic deletion of *Mboat7* promoted striking fatty liver and NAFLD progression but does not alter insulin sensitivity, suggesting the potential for cell autonomous roles. Here, we show that MBOAT7 function in adipocytes contributes to diet-induced metabolic disturbances including hyperinsulinemia and systemic insulin resistance. The expression of *Mboat7* in white adipose tissue closely correlates with diet-induced obesity across a panel of ∼100 inbred strains of mice fed a high fat/high sucrose diet. Moreover, adipocyte-specific genetic deletion of *Mboat7* is sufficient to promote hyperinsulinemia, systemic insulin resistance, and mild fatty liver. Unlike in the liver, where *Mboat7* plays a relatively minor role in maintaining arachidonic acid (AA)-containing PI pools, *Mboat7* is the major source of AA-containing PI pools in adipose tissue. Our data demonstrate that MBOAT7 is a critical regulator of adipose tissue PI homeostasis, and adipocyte MBOAT7-driven PI biosynthesis is closely linked to hyperinsulinemia and insulin resistance in mice.

## Introduction

The rising prevalence of obesity closely parallels the rise of nonalcoholic fatty liver disease (NAFLD), which is a leading cause of liver disease mortality in developed countries (Cohen et al., 2011; Wree et al., 2013; Rinella et al., 2016; Sarwar et al. 2018). Because of this rapidly growing health crisis, there is an increasing need for mechanistic insight to allow a path for the rational design of new therapeutic strategies. One important clue in NAFLD research has recently emerged where multiple genome wide association studies (GWAS) identified the common rs641738 single nucleotide polymorphism (SNP) located close to the lysophosphatidylinositol (LPI)-acylating enzyme, Membrane Bound O-Acyltransferase 7 (MBOAT7), as a risk allele. The rs641738 (C>T) SNP is associated with increased susceptibility to NAFLD (Mancina et al., 2016; Luukkonen et al., 2016; Teo et al., 2021), liver diseases of alcoholic and viral etiologies (Buch et al., 2015; Thabet et al., 2016; Thabet et al., 2017), and other complex metabolic diseases (Johansen et al., 2016; Jacher et al., 2019). Using an antisense oligonucleotide (ASO)-mediated approach we recently showed that *Mboat7* knockdown exacerbated high fat diet (HFD)-induced hepatic steatosis and inflammation, hyperinsulinemia, and insulin resistance (Helsley et al., 2019). It is important to note that we also showed that genetic deletion of the neighboring gene transmembrane channel like 4 (*Tmc4*) did not promote NAFLD (Helsley et al., 2019).

Very similar findings were reported by Meroni and colleagues using a morpholino oligonucleotide (MPO)-driven knockdown approach, further demonstrating that *Mboat7* loss of function promoted fatty liver, and importantly showed that *Mboat7* expression is suppressed by insulin (Meroni et al., 2020). More recently, several independent laboratories have shown that hepatocyte-specific deletion of *Mboat7* (*Mboat7*^HSKO^) worsens hepatic steatosis, inflammation, and fibrosis, but unlike the oligonucleotide-based knockdown, approaches does not impact insulin or glucose homeostasis (Tanaka et al., 2021; Thangapandi et al., 2021; Xia et al., 2021). Given the disparate results between the results of published ASO and MPO knockdown experiments (Helsley et al., 2019; Meroni et al., 2020) and those of these recent studies of *Mboat7*^HSKO^ (Tanaka et al., Thangapandi et al., 2021; Tanaka et al., 2021; Xia et al., 2021), we hypothesized that the function of MBOAT7 in adipocytes may be critically important in regulating circulating insulin levels and insulin action in target tissues. It is well appreciated that ASOs can effectively suppress target gene expression in adipocytes (Crooke et al., 2021; Helsley et al., 2019), and adipose tissue plays a critically important role in the progression of NAFLD and systemic glucose homeostasis. To better understand the tissue specific roles of MBOAT7, we generated adipocyte-specific (*Mboat7*^ASKO^) and hepatocyte-specific (*Mboat7*^HSKO^) knockout mice to understand the cell-autonomous contributions of *Mboat7* to HFD-induced metabolic disturbances. Here, we find that adipocyte *Mboat7* contributes less so to hepatic steatosis and injury when compared to hepatocyte *Mboat7*, but instead plays a critically important role in regulating local adipose tissue LPI/PI homeostasis, hyperinsulinemia, and systemic insulin sensitivity.

## Results

### Adipocyte-specific genetic deletion of *Mboat7* promotes mild fatty liver but profound hyperinsulinemia and insulin resistance

To understand the cell autonomous roles of *Mboat7* in HFD-driven metabolic disturbance we generated parallel colonies of either hepatocyte-specific (*Mboat7*^HSKO^) or adipocyte-specific (*Mboat7*^ASKO^) *Mboat7* knockout mice and subjected them to HFD feeding. Our initial rationale to study the role of *Mboat7* outside of the liver came from studies where we used a systems genetics approach to examine tissue-specific links between *Mboat7* expression and adiposity in mice represented in the hybrid mouse diversity panel (HMDP) when challenged with an obesity-promoting high fat and high sucrose diet (Ghazalpour et al., 2012; Parks et al., 2013). In our previous work, we found that *Mboat7* expression in the liver was only modestly correlated (r = - 0.244, p=0.01) with adiposity (Helsley et al., 2019). However, here we show that *Mboat7* expression in WAT is strongly negatively correlated with gonadal, subcutaneous, and retroperitoneal WAT in both female and male mice (**Figure 1a-1d**). It is interesting to note that WAT *Mboat7* mRNA expression is not significantly correlated with adiposity measures in mice maintained on a standard rodent chow (data not shown), suggesting a role for WAT *Mboat7* in selectively shaping diet-driven metabolic disturbance.

**Figure 1.**
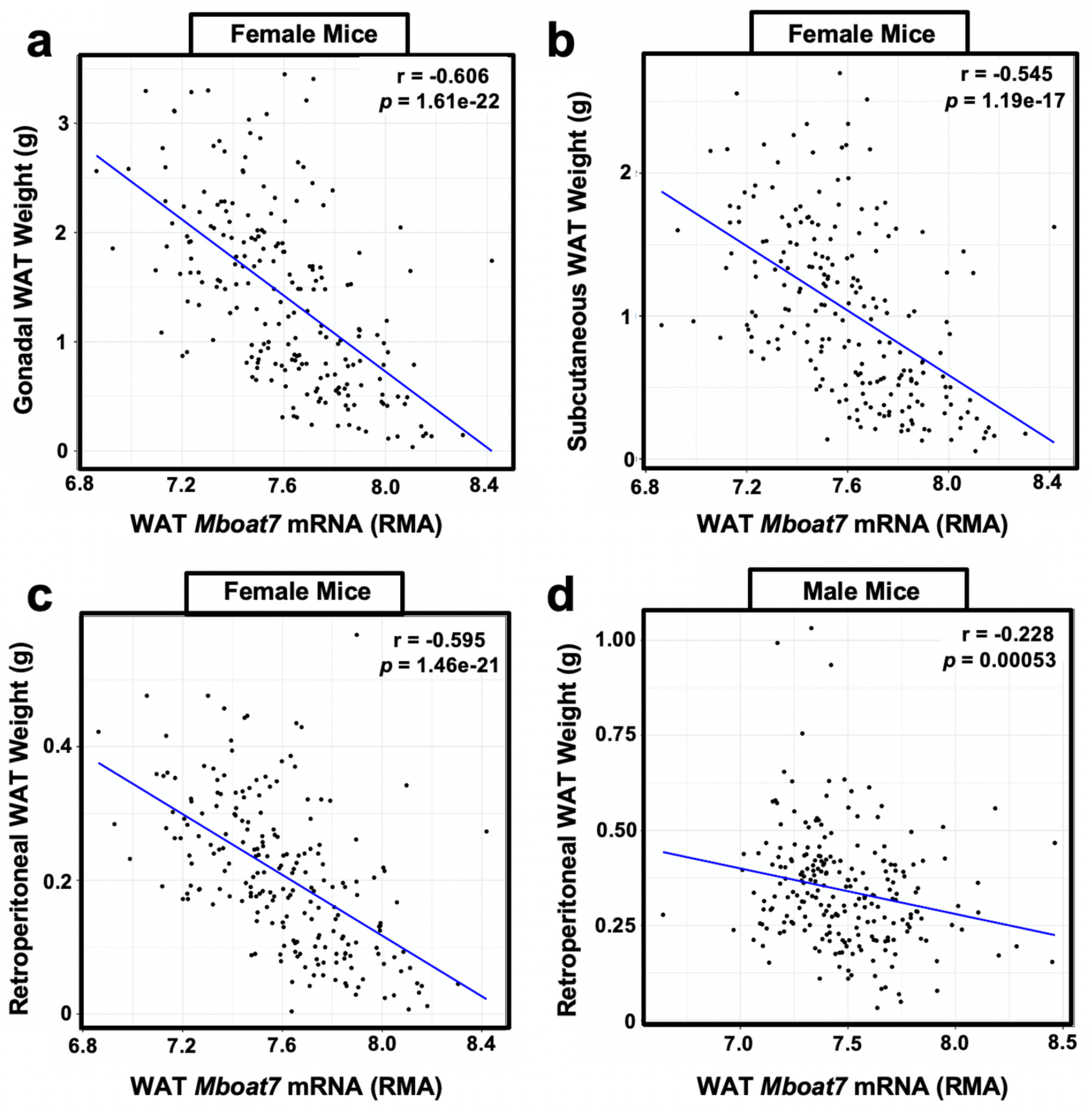
*Mboat7* Expression in White Adipose Tissue is Correlated with Adiposity in mice. We used a systems genetics approach to examine links between *Mboat7* expression and metabolic traits in mice from the hybrid mouse diversity panel (HMDP). To induce obesity, all mouse strains represented in the HMDP were fed an obesity-promoting high fat and high sucrose diet. Across the different strains in the HMDP, the expression (RMA, Robust Multi-Array Average) of *Mboat7* in adipose tissue has a strong negatively correlated with gonadal (**a**), subcutaneous (**b**), and retroperitoneal (**c**) white adipose tissue mass in females. There is a similar negative correlation in male (**d**) retroperitoneal adipose tissue.

To generate congenic adipocyte-specific *Mboat7* knockout mice (*Mboat7*^ASKO^) we crossed mice harboring a post-FLP recombinase conditionally-targeted *Mboat7* floxed allele (Anderson et al., 2013) to mice transgenically expressing Cre recombinase under the adiponectin promoter/enhancer (Eguchi et al., 2011), and then backcrossed these mice > 10 generations into the C57BL/6J background. Compared to control mice (*Mboat7*^flox/flox^), *Mboat7*^ASKO^ mice had significantly reduced MBOAT7 protein expression in gonadal WAT, subcutaneous WAT, and subscapular brown adipose tissue (BAT) and a modest increase in skeletal muscle (**Figure 2a**). It is important to note that liver MBOAT7 protein levels were not different in *Mboat7*^ASKO^ mice (**Figure 2a**). However, the molecular weight of the major MBOAT7 protein isoform in the liver is slightly smaller when compared to the major isoform in WAT and BAT (**Figure 2a**). Adipocyte-specific deletion of *Mboat7* promotes marked reorganization of MBOAT7 substrate (LPI) and product (PI) lipids in gonadal WAT (**Figure 2b-d**). Particularly when challenged with a HFD, *Mboat7*^ASKO^ mice have significantly elevated levels of 16:0 LPI, 18:0 LPI, and 18:1 LPI lipids in gonadal WAT (**Figure. 2b**). Reciprocally, the predominant enzymatic product of MBOAT7 (38:4 PI) is reduced by > 60% in *Mboat7*^ASKO^ mice under both chow and HFD feeding conditions (**Figure 2c**). Although less abundant, 36:4 PI and 38:5 PI lipids are also reduced in the gonadal WAT of *Mboat7*^ASKO^ mice, and there is a reciprocal increase in other more saturated species of PI (34:2 PI, 36:2 PI, and 36:3 PI) (**Figure 2d**). It is important to note that the levels of LPI and PI lipids in the plasma and liver were unaffected in *Mboat7*^ASKO^ mice (**Figure 2 – figure supplement 1**), which is in contrast to what has been recently reported in *Mboat7*^HSKO^ mice where both liver and plasma LPI and PI homeostasis is altered (Tanaka et al., 2021; Thangapandi et al., 2021; Xia et al., 2021). Collectively, these results demonstrate that MBOAT7 is a major regulator of the local LPI and PI lipidome in gonadal WAT, but unlike *Mboat7*^HSKO^ mice, adipocyte-specific deletion of *Mboat7* does not significantly alter circulating or liver LPI/PI homeostasis.

**Figure 2.**
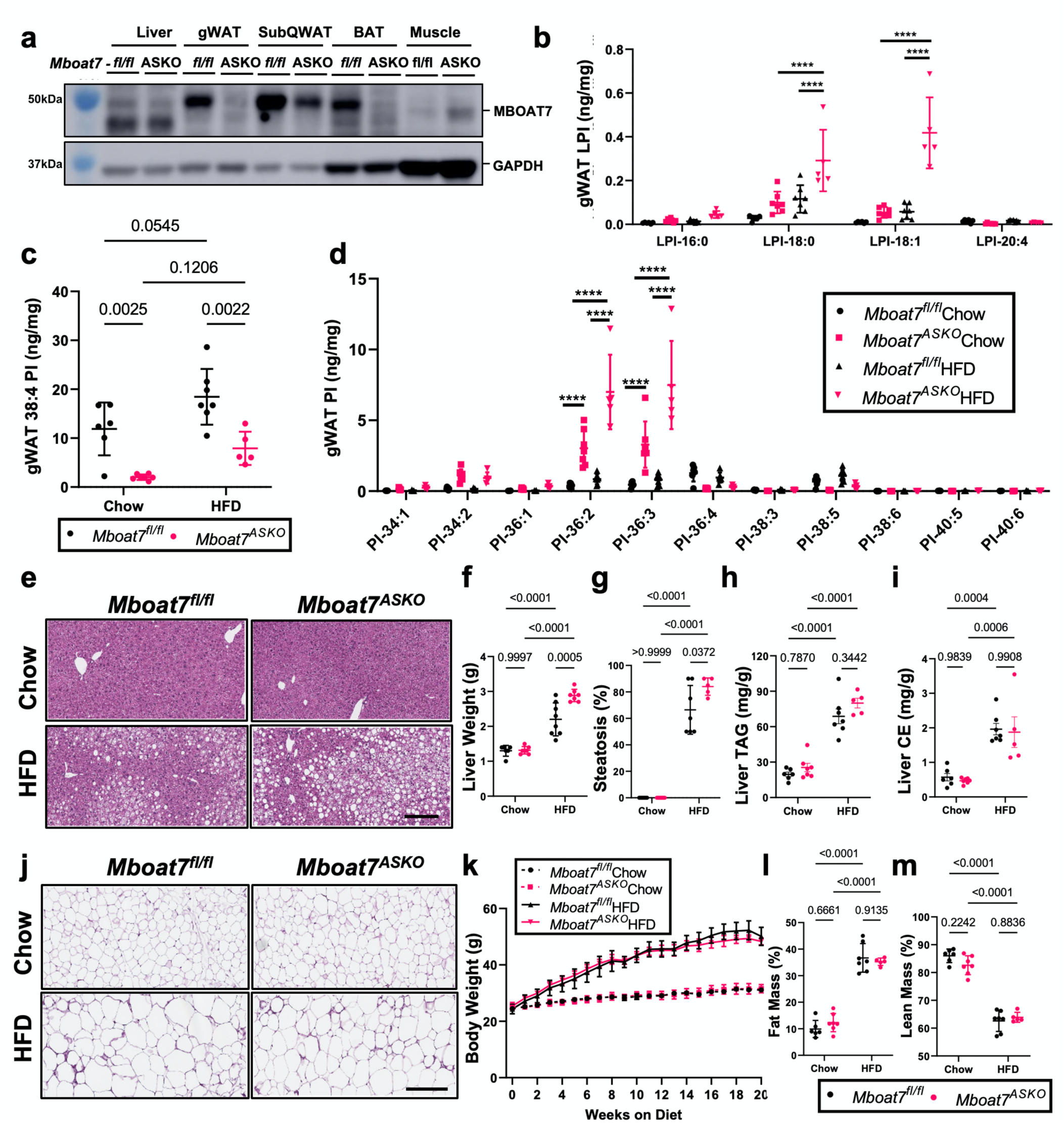
Adipocyte-Specific *Mboat7* Deletion (*Mboat7^ASKO^*) Promotes Mild Fatty Liver. Male control (*Mboat7^fl/fl^*) or adipocyte-specific Mboat7 knockout mice (*Mboat7^ASKO^*) were fed chow or high fat diet (HFD) for 20-weeks and metabolically phenotyped. (**a**) Western blots from tissues collected from *Mboat7^fl/fl^* or *Mboat7^ASKO^*mice. (**b-n**) Gonadal white adipose tissue (gWAT) lysophosphatidylinositol (LPI) (**b**) and phosphatidylinositol (PI) species, including the MBOAT7 product PI-38:4 (**c**) and others (**d**), were quantified via liquid chromatography-tandem mass spectrometry in *Mboat7^fl/fl^* or *Mboat7^ASKO^* mice were fed chow or high fat diet (HFD) for 20-weeks (n=5-7; \*\*\*\**P*≤0.0001; Two-way (**c**) or Three-way (**b,d**) ANOVA with Tukey’s *post-hoc* test). (**e**) Representative liver hematoxylin and eosin stained sections. 10x magnification (scale bar=200µm). (**f**) Liver weight measurements from *Mboat7^fl/fl^* or *Mboat7^ASKO^*mice fed Chow and HFD for 20 weeks (n=6-8; Two-way ANOVA with Tukey’s *post-hoc* test). (**g**) Percent steatosis was quantified by a blinded pathologist (n=5-7; Two-way ANOVA with Tukey’s *post-hoc* test). Hepatic triglycerides (**h**) and hepatic esterified cholesterol (**i**) were measured enzymatically (n=5-7; Two-way ANOVA with Tukey’s *post-hoc* test). (**j**) Representative gWAT hematoxylin and eosin stained sections. 10x magnification (scale bar=200µm). (**k**) Body weight was measured weekly. % Fat (**l**) and % Lean (**m**) mass were determined via echo-MRI after 8 weeks of chow or HFD in *Mboat7^fl/fl^* or *Mboat7^ASKO^*mice (n=5-7; Two-way ANOVA with Tukey’s *post-hoc* test). All data are presented as mean ± S.D.

Next, we compared and contrasted HFD-driven metabolic phenotypes in adipocyte-specific (*Mboat7*^ASKO^) and hepatocyte-specific (*Mboat7*^HSKO^) mice. In agreement with what has been previously reported by three independent groups (Tanaka et al., 2021; Thangapandi et al., 2021; Xia et al., 2021), we also find that *Mboat7*^HSKO^ mice have significant reductions in 38:4 PI and elevation in its substrate LPI in the liver, but not in gonadal WAT (**Figure 2 – figure supplement 2g-2l**). *Mboat7*^HSKO^ mice have profound hepatic steatosis and elevated alanine aminotransferase (ALT) (**Figure 2 – figure supplement 2c-2f**), confirming the concept that MBOAT7 activity in hepatocytes opposes hepatic steatosis and liver injury (Tanaka et al., 2021; Thangapandi et al., 2021; Xia et al., 2021). Interestingly, adipocyte-specific deletion of *Mboat7* also promotes mild hepatomegaly and hepatic steatosis (**Figure 2e-2g**). However, the significantly increased hepatic triglyceride and cholesterol ester levels apparent in *Mboat7*^HSKO^ (Tanaka et al., 2021; Thangapandi et al., 2021; Xia et al., 2021) are not seen in *Mboat7*^ASKO^ mice (**Figure 2h,2i**). It is important to note that upon HFD feeding total body weight, fat mass, lean mass, and WAT histology are similar in *Mboat7*^HSKO^ when compared to their controls (**Figure 2 – figure supplement 2b**; and data not shown) as well as *Mboat7*^ASKO^ compared to their appropriate controls (**Figure 2j-2m**). Collectively, these data suggest that MBOAT7 has clear cell autonomous roles in regulating circulating and tissue LPI and PI pools and MBOAT7-driven LPI acylation in hepatocytes is the primary driver of hepatic steatosis seen with MBOAT7 loss of function.

### Adipocyte-specific, but not hepatocyte-specific, deletion of *Mboat7* promotes hyperinsulinemia and insulin resistance

Obesity and T2DM are commonly seen in subjects with NAFLD, and it is well appreciated that excessive accumulation of lipids (i.e. “lipotoxicity) in the liver, skeletal muscle, and pancreatic beta cells can drive the overproduction of insulin and insulin resistance in target tissues (Unger 1995; Unger et al., 2010; Samuel et al., 2018). Recent studies have demonstrated that the hepatic expression of MBOAT7 is suppressed in obese humans and rodents (Helsley et al., Meroni et al. 2020), and insulin treatment can acutely suppress MBOAT7 mRNA and protein expression (Meroni et al., 2020). Furthermore, we demonstrated that ASO-mediated knockdown of *Mboat7* in HFD-fed mice elicits profound hyperinsulinemia and systemic insulin resistance (Helsley et al., 2019). However, several recent studies have demonstrated that genetic deletion of *Mboat7* specifically in hepatocytes (i.e. *Mboat7*^HSKO^ mice) does not alter insulin action or glucose tolerance (Tanaka et al., 2021; Xia et al., 2021). To further understand the cell autonomous roles of *Mboat7* in insulin production and insulin action, we performed a series of studies comparing adipocyte-specific (*Mboat7*^ASKO^) and hepatocyte-specific (*Mboat7*^HSKO^) *Mboat7* knockout mice (**Figure 3; Figure 2 – figure supplement 2m-2p**). Despite having a severe fatty liver (**Figure 2 – figure supplement 2i-2k**), *Mboat7*^HSKO^ mice have normal glucose tolerance when challenged with a HFD (**Figure 2 – figure supplement 2m).** HFD-fed *Mboat7*^HSKO^ mice also have similar levels of fasting blood glucose, insulin, and leptin when compared to *Mboat7*^flox/flox^ controls (**Figure 2 – figure supplement 2n-2p**). In contrast, when challenged with a HFD *Mboat7*^ASKO^ mice exhibit impaired systemic glucose tolerance and increased fasting blood glucose and insulin levels (**Figure 3a-3d**). It is important to note that most metabolic studies were performed in male *Mboat7*^ASKO^ mice, but many of the lipid changes and glucose intolerance phenotypes were also apparent in female *Mboat7*^ASKO^ mice (**Figure 3 – figure supplement 1**). When we subjected *Mboat7*^ASKO^ mice to an insulin tolerance test (ITT) to assess *in vivo* insulin sensitivity, there were some apparent differences between male and female mice **(Figure 3 – figure supplement 2)**. When maintained on a chow diet, both male and female *Mboat7*^ASKO^ mice had similar ITT responses to those seen in *Mboat7*^flox/flox^ control mice **(Figure 3 – figure supplement 2a,2b,2c,2d)**. In contrast, when challenged with a high fat diet, male *Mboat7*^ASKO^ mice had an unexpected increase in blood glucose levels, whereas HFD-fed *Mboat7*^flox/flox^ control mice maintained glucose levels after an insulin challenge indicating clear HFD-induced insulin resistance (**Figure 3 – figure supplement 2e,2f**). Whereas HFD-fed female *Mboat7*^flox/flox^ control mice showed appreciable insulin-induced lowering of blood glucose, HFD-fed *Mboat7*^ASKO^ mice showed significantly blunted insulin action (**Figure 3 – figure supplement 2g,2h**). It is interesting to note that gonadal WAT tissue levels of 18:0- and 18:1-containing LPIs were significantly correlated with fasting blood glucose and GTT area under the curve (**Figure 3 – figure supplement 3**). Collectively, these data show for the first time that MBOAT7-driven LPI acylation in adipocytes, but not in hepatocytes, is critically important for maintenance of circulating insulin and glucose levels as well as systemic insulin action.

**Figure 3.**
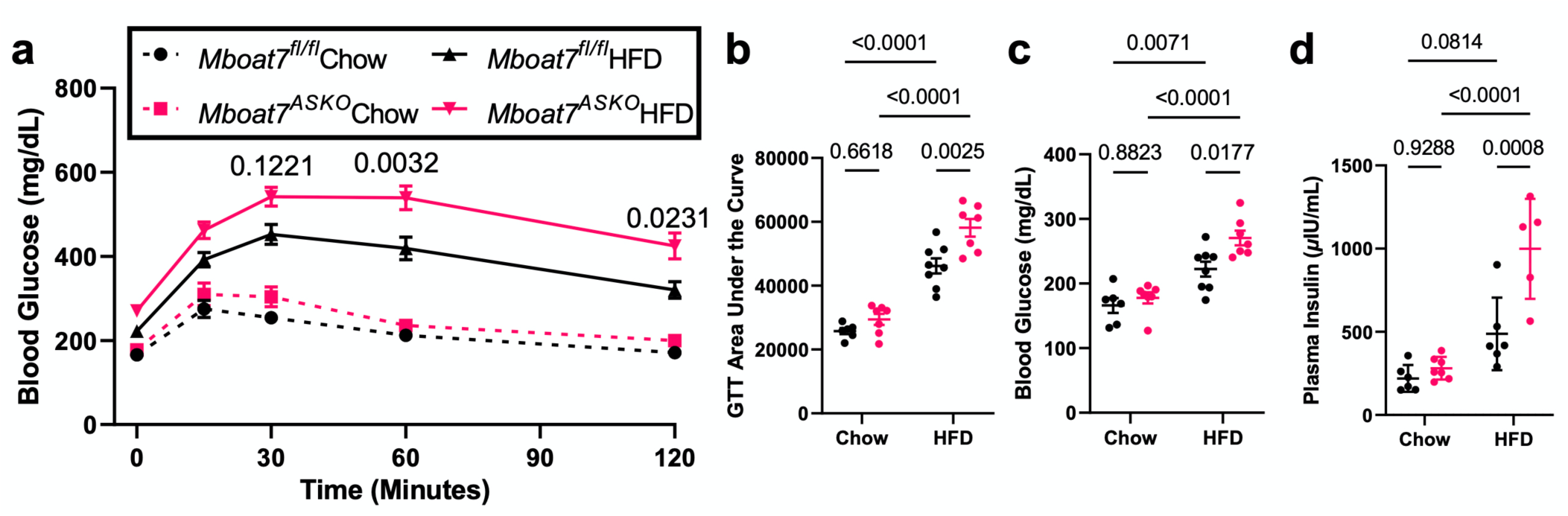
Adipocyte-Specific *Mboat7* Deletion (*Mboat7^ASKO^*) Promotes Hyperinsulinemia and Glucose Intolerance. (**a-c**) Male control (*Mboat7^fl/fl^*) or adipocyte-specific Mboat7 knockout mice (*Mboat7^ASKO^*) were fed a chow or HFD for 12 weeks and then underwent an intraperitoneal glucose tolerance test (GTT). (**a**) Plasma glucose levels were measured (in duplicate or triplicate at each timepoint) throughout the GTT (n=5-7; Three-way ANOVA with Tukey’s *post-hoc* test). (**b**) The area under the curve was calculated for each mouse throughout the GTT (n=5-7; Two-way ANOVA with Tukey’s *post-hoc* test). (**c**). Fasting blood glucose was measured after a four hour fast (time=0 minutes for GTT) (n=5-7; Two-way ANOVA with Tukey’s *post-hoc* test). (**d**) Fasting plasma insulin was measured in *Mboat7^fl/fl^*or *Mboat7^ASKO^* mice that were fed a chow or HFD for 20 weeks (n=5-7; Two-way ANOVA with Tukey’s *post-hoc* test).

### *Mboat7* is a critical metabolic regulator of adipose tissue lipid homeostasis and adipose tissue function

The proper storage of excess energy in the form of triacylglycerol (TAG) within adipose tissue is critically important for overall metabolic health. Although the major lipid class stored in WAT is TAG, adipocytes play a critically important role in regulating the levels of other lower abundance lipids that can shape local and systemic inflammatory and hormonal responses to influence T2DM, NAFLD, and other related metabolic diseases. To further understand how MBOAT7 impacts adipose tissue metabolism, we performed a series of comprehensive lipidomics studies in gonadal WAT isolated from chow- and HFD-fed *Mboat7*^flox/flox^ control mice and *Mboat7*^ASKO^ mice. Although MBOAT7 has been previously shown to have exquisite substrate selectively toward saturated LPIs and arachidonyl-CoA (Gijón et al., 2008; Lee et al., 2012; Zarini et al., 2014), we also analyzed the abundance of other free fatty acids, oxylipins, and diverse species of neutral lipids and glycerophospholipids. First, as a reference, it is notable that hepatocyte-specific deletion of *Mboat7* (*Mboat7*^HSKO^ mice) results in a large accumulation of substrate LPIs and a modest reduction in 38:4 PI in the liver, but this is not apparent in gonadal WAT (**Figure 2 – figure supplement 2g-2l**). In contrast, *Mboat7*^ASKO^ mice do not show alteration in LPI or PI lipids in the liver (**Figure 2 – figure supplement 1d-1f**) but do have elevations in 18:0- and 18:1- containing LPIs and large reductions in 38:4 PI in gonadal WAT (**Figure 2b-2d**). *Mboat7*^ASKO^ mice also have reciprocal increases in more saturated PI species (34:2, 36:2, and 36:3) compared to *Mboat7*^flox/flox^ control mice in gonadal WAT (**Figure 2d**), but not liver (**Figure 2 – figure supplement 1f**). These results demonstrate that MBOAT7 plays a critical role in maintaining LPI and PI levels within the local tissue context, and shows that MBOAT7 is the major enzymatic source of 38:4 in WAT (**Figure 2c**), but a more minor contributor in the liver (**Figure 2 – figure supplement 2h**; Tanaka et al., 2021; Thangapandi et al., 2021; Xia et al., 2021).

Given the critical role that MBOAT7 plays in WAT LPI/PI homeostasis, we wanted to more comprehensively understand whether other lipid classes were altered in the WAT of *Mboat7*^ASKO^ mice. Other than modest alterations in 36:2 and 36:3 phosphatidylethanolamine (PE), adipocyte-specific deletion of *Mboat7* did not result in significant alterations of free fatty acids, non-LPI lysophospholipids, phosphatidylcholines (PC), sphingomyelins (SM), or ceramides (CM) (**Figure 2 – figure supplement 3**). Given the critical role that MBOAT7 plays in the Land’s cycle remodeling pathway (Valentine et al., 2020), we also wanted to analyze arachidonic acid-derived oxylipins. When we quantified the WAT levels of > 60 species of oxylipins, none were significantly different between *Mboat7*^flox/flox^ control mice and *Mboat7*^ASKO^ mice (**Figure 2 – figure supplement 4**). Also, when we quantified the levels of neutral lipids such as diacylglycerols (DAG) and triacylglycerols (TAG) there were no significant differences between *Mboat7*^flox/flox^ control mice and *Mboat7*^ASKO^ mice when fed a HFD, but select species of TAG (52:3, 56:4, 50:5, etc.) were modestly reduced in chow-fed *Mboat7*^ASKO^ mice (**Figure 2 – figure supplement 5**). Despite the conservation seen in the major lipid classes found in WAT, *Mboat7*^ASKO^ mice did have significant alterations in some complex lipids containing either arachidonic acid (AA, 20:4, n-6) or eicosapentaenoic acid (EPA, 20:5, n-3) in WAT (**Figure 2 – figure supplement 6**). Specifically, some AA-containing PE species were increased in the WAT from *Mboat7*^ASKO^ mice (**Figure 2 – figure supplement 6a**). Furthermore, EPA-containing PI (38:5 PI) was dramatically reduced in *Mboat7*^ASKO^ mice, while other EPA-containing PS and PE species were slightly elevated (**Figure 2 – figure supplement 6c**). These data support the notion that adipocyte MBOAT7 plays a central role in WAT glycerophospholipid homeostasis locally but does not appreciably impact LPI and PI balance in the circulation of the liver.

Given the alterations in WAT lipid homeostasis in *Mboat7*^ASKO^ mice, we wanted to examine key aspects of adipose tissue function and cellularity within the adipose organ. Although body weight and total fat mass were not significantly different between *Mboat7*^flox/flox^ control mice and *Mboat7*^ASKO^ mice after 8 weeks on diet (**Figure 2k,2l**), after 20 weeks of HFD feeding the weight of gonadal WAT was significantly lower in *Mboat7*^ASKO^ mice (**Figure 4a**). In parallel, circulating levels of the adipocyte-derived hormone leptin were also reduced in HFD-fed *Mboat7*^ASKO^ mice (**Figure 4b**). Next, we wanted to examine whether basal or catecholamine-stimulated lipolysis (i.e. a critical physiological function of WAT) was altered in adipocyte-specific *Mboat7* knockout mice. Although in general the circulating levels of lipolysis product glycerol and non-esterified fatty acids (NEFA) were not altered under basal conditions or stimulated conditions in *Mboat7* deficient mice, chow-fed *Mboat7*^ASKO^ mice did have elevations in circulating glycerol when treated with the β3-adrenergic receptor agonist CL-326,243 (**Figure 4c,4d**). Next, we performed bulk RNA sequencing in gonadal WAT, and under both chow and HFD feeding conditions *Mboat7*^ASKO^ mice exhibited differential gene expression linked to altered lipid metabolism and immune cell populations (**Figure 4e,4f,4g**). In particular, *Mboat7*^ASKO^ mice had elevated WAT mRNA expression of genes primarily expressed in macrophages including a cluster of differentiation 68 (*Cd68*), integrin alpha X (*Itgax*, encoding CD11c), c-type lectin domain containing 7a (*Clec7a*), among others (**Figure 4f,4g**) indicating the potential for an altered abundance of adipose tissue macrophages. To further understand whether the abundance of macrophages and other immune cell populations were altered in *Mboat7*^ASKO^ mice we isolated the stromal vascular fraction from gonadal WAT and performed flow cytometric analysis of immune cell populations (**Figure 4h-4j & Figure 4 – figure supplement 1**). Of the adipose tissue macrophage populations identified, *Mboat7*^ASKO^ mice had elevated levels of Cd11c-/Cd206+ cell populations and a reciprocal decrease in Cd11c+/Cd206+ double-positive cells (**Figure 4g**). Although the relative percentage of Cd11c-/Cd206+ cells was not significantly correlated with body weight (Figure 4h), the Cd11c+/Cd206+ double-positive cell population was significantly positively correlated with body weight in this cohort (**Figure 4i,4j**). In addition to alterations in macrophage subsets, HFD-fed *Mboat7*^ASKO^ mice had elevated T cells and B cells compared to control (*Mboat7*^flox/flox^) mice **(Figure 4 – figure supplement 1b-1f)**. These data demonstrate that adipocyte MBOAT7-driven LPI acylation plays an important role in WAT lipid metabolic and immune cell homeostasis which can then shape systemic glucose tolerance.

**Figure 4.**
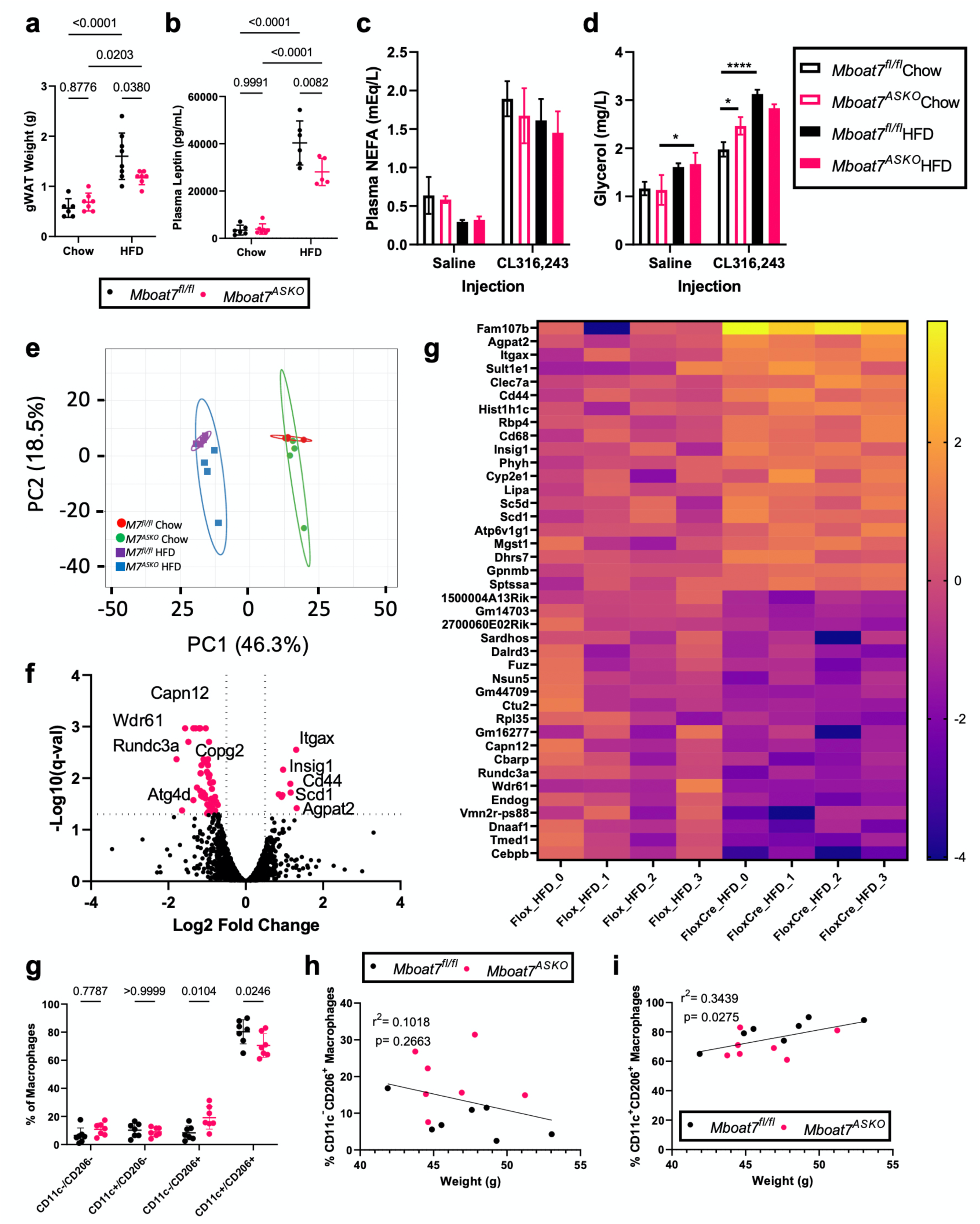
Adipocyte-Specific *Mboat7* Deletion (*Mboat7^ASKO^*) Reorganizes White Adipose Tissue Gene Expression and Circulating Adipokine Levels. Male control (*Mboat7^fl/fl^*) or adipocyte-specific Mboat7 knockout mice (*Mboat7^ASKO^*) were fed chow or high fat diet (HFD) for 20-weeks. (**a**) gWAT weight measurements from *Mboat7^fl/fl^* or *Mboat7^ASKO^* mice fed Chow and HFD for 20 weeks (n=6-8; Two-way ANOVA with Tukey’s *post-hoc* test). (**b**) Plasma Leptin was measured in *Mboat7^fl/fl^* or *Mboat7^ASKO^* mice that were fed a chow or HFD for 20 weeks (n=5-7; Two-way ANOVA with Tukey’s *post-hoc* test). To assess β3-adrenergic stimulated lipolysis, plasma non-esterified fatty acids (NEFA) (**c**) and glycerol (**d**) were measured in *Mboat7^fl/fl^* or *Mboat7^ASKO^* mice fed a chow or HFD for 11 weeks 15 minutes after saline or CL316,243 injection (n=2-5; Three-way ANOVA with Tukey’s *post-hoc* test). (**e-g**) gWAT RNA was used for RNA-sequencing from *Mboat7^fl/fl^*or *Mboat7^ASKO^* mice that were fed a chow or HFD for 20 weeks (**e**) Groups clustered primarily based on the diet by principal component analysis (n=4/group). (**f**) A Volcano plot of transcripts was used to determine differentially expressed genes (DEGs) in *Mboat7^fl/fl^* or *Mboat7^ASKO^*mice that were fed an HFD for 20 weeks. Plot summarizes log2 fold changes vs significance in response to *Mboat7* inhibition (n=4; genes with q-val < 0.05 and fold change > |0.5| were considered significantly differentially expressed). (**g**) Row-normalized expression for the top 20 up and downregulated DEGs are shown by heat map in *Mboat7^fl/fl^* or *Mboat7^ASKO^* mice that were fed an HFD for 20 weeks. **(h)** gWAT stromal vascular fraction was subjected to flow cytometry analysis of macrophage subpopulations. **(I,j)** correlation between gWAT macrophage subsets and body weight. All data are presented as mean ± S.D. unless otherwise noted.

## Discussion

Since the original GWAS study by Buch and colleagues linking the rs641738 SNP near *MBOAT7* to liver disease (Buch et al., 2015), there has been rapid progress in our understanding of how *MBOAT7* is mechanistically linked to the progression of alcohol-associated liver disease (ALD), NAFLD, and viral-driven liver injury. The clear association between MBOAT7 loss of function and diverse liver diseases serves as yet another example of how genetics can powerfully identify new pathways relevant to human disease. It also supports the long-standing notion that abnormal lipid metabolism initiates liver injury. Since 2019, several animal studies have likewise demonstrated that *Mboat7* loss of function in mice is sufficient to drive NAFLD progression (Helsley et al., 2019; Meroni et al., 2020; Tanaka et al., 2021; Thangapandi et al., 2021; Xia et al., 2021). This manuscript builds on our initial observation that ASO-mediated knockdown of *Mboat7* promotes NAFLD progression, hyperinsulinemia, and insulin resistance in mice (Helsley et al., 2019). Here we have further clarified the cell autonomous roles of *Mboat7* in HFD-driven metabolic disturbance by comparing metabolic phenotypes in adipocyte-specific (*Mboat7*^ASKO^) and hepatocyte-specific (*Mboat7*^HSKO^) mice. The major findings of the current study include the following: (1) *Mboat7* expression in WAT is negatively correlated with adiposity across the strains represented in the Hybrid Mouse Diversity Panel, (2) Hepatocyte-specific deletion of *Mboat7* (*Mboat7*^HSKO^) promotes fatty liver and liver injury, but does not alter circulating insulin levels or tissue insulin sensitivity in HFD-fed mice, (3) Adipocyte-specific deletion of *Mboat7* (*Mboat7*^ASKO^) promotes mild fatty liver, and clear hyperinsulinemia and systemic insulin resistance in HFD-fed mice, (4) MBOAT7 is the major source of arachidonic acid containing PI (38:4 PI), and also indirectly impacts other glycerophospholipids in gonadal WAT, (5) MBOAT7 function in gonadal WAT does not alter the tissue abundance of arachidonic acid-derived oxylipins, and (6) MBOAT7 function in gonadal WAT suppresses adipose tissue weight, circulating leptin levels, and impacts the immune cell landscape in WAT. Collectively, these data support a clear cell autonomous role for MBOAT7-driven acylation of LPI lipids as a key protective mechanism against obesity-linked NAFLD progression, hyperinsulinemia, and systemic insulin resistance. These data strongly suggest that MBOAT7 is an important contributor to multiple aspects of the metabolic syndrome, which is regulated by the combination of MBOAT7 function in hepatocytes, adipocytes, and likely other cell types that contribute to tissue inflammation and fibrosis.

One of the key findings of this work is that MBOAT7 function in adipocytes is critically important for the maintenance of euglycemia in obese mice (**Figure 3**). In contrast, genetic deletion of *Mboat7* in hepatocytes is sufficient to drive hepatic steatosis but does not alter hyperinsulinemia or insulin sensitivity in either chow or HFD-fed mice (**Figure 2 – figure supplement 2**). These new clues now raise the question of how MBOAT7 in adipocyte controls insulin secretion in the pancreas or insulin sensitivity in the liver or skeletal muscle which are the predominant sites of maintaining systemic euglycemia. The most straightforward explanation is that alterations in either the substrates (LPIs or fatty acyl-CoAs) or products (38:4 PI) of the MBOAT7 reaction initiate some local or endocrine lipid signaling effects. We originally hypothesized that when adipocyte MBOAT7 function is lost, that substrate LPIs would accumulate in both the WAT as well as the circulation to initiate a WAT-to-pancreas endocrine axis to stimulate insulin overproduction. In support of this concept, several published reports are showing that LPI lipids can stimulate glucose-stimulated insulin secretion (GSIS) in pancreatic beta cells in culture (Metz SA, 1986; Liu et al.,2016). Furthermore, 18:1 LPI has also been shown to stimulate the key metabolic hormone glucagon-like peptide 1 (GLP-1) in intestinal enteroendocrine *L*-cells to further shape insulin secretion (Arifin et al., 2018; Harada K., et al. 2017). However, our results suggest that although WAT tissue of LPI is dramatically increased in *Mboat7*^ASKO^ mice (Figure 2b), circulating levels of LPI lipids are unaltered in *Mboat7*^ASKO^ mice (**Figure 2 – figure supplement 1**). These data suggest this potential endocrine axis is unlikely, and instead an autocrine or paracrine axis is more likely within WAT. It is important to note that LPI lipids can exhibit altered signaling potential when bound to albumin versus being carried on plasma lipoproteins (Kurano et al., 2021), so additional work is required to determine whether adipocyte MBOAT7 activity selectively impacts albumin-conjugated versus lipoprotein-associated LPI levels. In conclusion, this work shows that MBOAT7 function in adipocytes and hepatocytes play unique roles in shaping HFD-driven metabolic disturbance, and further supports the notion that the LPI-MBOAT-PI axis may have untapped therapeutic potential in obesity-related insulin resistance and NAFLD progression.

## METHODS

### Key Resources Table

| Key Resource Table |  |  |  |  |  |
| --- | --- | --- | --- | --- | --- |
| Reagent type or resource | Designation | Source or Reference | Identifiers | Additional Information | Research Resource Identifiers (RRID) |
| Genetic reagent (*M.musculus*) | *Mboat7tm1a(KOMP)Wtsi/Mboat7tm1a(KOMP)Wtsi* | PMID: 23472195 |  |  | RRID: MGI:5510874 |
| Genetic reagent (*M.musculus*) | *B6.FVB-Tg(Adipoq-cre)1Evdr/J* | Jackson Laboratory | Stock#: 028020 |  | RRID: IMSR_JAX:028020 |
| Genetic reagent (*M.musculus*) | *B6.Cg-Speer6-ps1Tg(Alb-cre)21Mgn/J* | Jackson Laboratory | Stock#: 003574 |  | RRID: IMSR_JAX:003574 |
| Antibody | Anti-MBOAT7 (Rat monoclonal) | PMID: 23097495 |  | 1:1000 | RRID: AB_2813851 |
| Antibody | Anti-Rat-HRP | Cell Signaling | Cat#: 7077 | 1:5000 | RRID: AB_10694715 |
| Antibody | Anti-GAPDH-HRP (Rabbit monoclonal) | Cell Signaling | Cat#: 8884 | 1:5000 | RRID: AB_11129865 |
| Antibody | BV510 Rat Anti-Mouse CD45 | BD Pharm | Cat#: 563891 | 1:100 | RRID: AB_2734134 |
| Antibody | Anti-CD3e-APC-Cy7 (Hamster monoclonal) | BD Pharm | Cat#: 557596 | 1:100 | RRID: AB_396759 |
| Antibody | Ly-6G/Ly-6C Monoclonal Antibody (RB6-8C5), FITC | Thermo | Cat#: 11-5931-82 | 1:100 | RRID: AB_465314 |
| Antibody | CD45R (B220) Monoclonal Antibody (RA3-6B2), eFluor™ 506 | eBioscience | Cat#: 69-0452-82 | 1:100 | RRID: AB_2637455 |
| Antibody | Anti-Cd11b-PE-Cy7 (Rat monoclonal) | BD Pharm | Cat#: 552850 | 1:100 | RRID: AB_394491 |
| Antibody | PerCP/Cyanine5.5 anti-mouse F4/80 Antibody | Biolegend | Cat#:123128 | 1:100 | RRID: AB_893484 |
| Antibody | CD115 (c-fms) Monoclonal Antibody (AFS98), PE | eBioscience | Cat#: 12-1152-83 | 1:100 | RRID: AB_465809 |
| Antibody | Anti-Cd11c-Alexa Fluor 700-(Hamster monoclonal) | BD Pharm | Cat#: 560583 | 1:100 | RRID: AB_1727421 |
| Antibody | Anti-CD206 Alexa Fluor 647 (Rat monoclonal) | Biolegend | Cat#: 141712; | 1:100 | RRID: AB_10900420 |
| Commercial assay kit | ALT kit | Sekisui Diagnostics | 318-30 |  |  |
| Commercial assay kit | Liver TG | Wako | 994-02891 |  |  |
| Commercial assay kit | Free Cholesterol | Wako | 993-02501 |  |  |
| Commercial assay kit | Total Cholesterol | Fisher Sci | TR134321 |  |  |
| Commercial assay kit | Metabolic Hormone Combo 1 | MesoScale Discovery | Cat#: K15306K |  |  |
| Commercial assay kit | NEFA Kit | Wako | 999-34691, 995-34791, 991-34891, 993-35191, 276-76491 |  |  |
| Commercial assay kit | Glycerol kit |  |  |  |  |
| Commercial assay kit | C-Peptide ELISA | Crystal Chem | 90050 |  |  |
| Chemical compound, drug | Beta-3 AR Agonist | Sigma | C5976-5mg |  |  |
| Chemical compound, drug | Glucose | Sigma | G8270 |  |  |
| Chemical compound, drug | Insulin (HumalinR) | Lily | 0002-8215-17 |  |  |

### Mice and experimental diets

To generate conditional *Mboat7* knockout mice, we obtained “knockout first” (Mboat7^tm1a(KOMP)Wtsi^) mice from Dr. Philip Hawkins (Anderson et al., 2013), and crossed these mice with mice transgenically expressing FLP recombinase to remove the NEO cassette resulting in a conditional *Mboat7* floxed allele. The FLP transgene was then subsequently bred out of the line and resulting *Mboat7*^flox/WT^ mice were then either bred with transgenic mice expressing CRE recombinase under either the adiponectin promoter/enhancer (for adipocyte-specific deletion) or under the albumin promoter (for hepatocyte-specific deletion). Importantly, each substrain of mice was backcrossed these mice > 10 generations into the C57BL/6J background to produce congenic lines, and confirmation of sufficient backcrossing into the C57BL/6J background was confirmed by mouse genome single nucleotide polymorphism (SNP) scanning at the Jackson Laboratory (Bar Harbor, ME). Age-matched animals were put on either standard rodent chow (Teklad TD:130104) or 60% kcal fat high fat diet (HFD) (Research Diets D12492) for up to 20 weeks. Glucose and insulin tolerance tests were performed as previously described (Schugar et al., 2017; Gromovsky et al., 2017; Warrier et al., 2015; Brown et al., 2010; Brown et al., 2008a; Brown et al. 2008b) in mice following 12 and 14 weeks of concurrent chow and HFD-feeding, respectively. Quantitation of lean and fat mass were done using an EchoMRITM-130 Body Composition Analyzer (EchoMRI International). All mice were maintained in an Association for the Assessment and Accreditation of Laboratory Animal Care, International-approved animal facility, and all experimental protocols were approved by the Institutional Animal Care and Use Committee of the Cleveland Clinic (IACUC protocols # 2018-2053 and # 00002499).

### Standardized necropsy conditions

To keep results consistent, the experimental mice were fasted for 4 hours (from 09:00 to 13:00) prior to necropsy. At necropsy, all mice were terminally anesthetized with ketamine/xylazine (100-160mg/kg ketamine-20-32mg/kg xylazine), and a midline laparotomy was performed. Blood was collected by heart puncture. Following blood collection, a whole-body perfusion was conducted by puncturing the right atria and slowly delivering 10 ml of saline into the left ventricle of the heart to remove blood from tissues. Tissues were collected and immediately snap frozen in liquid nitrogen for subsequent biochemical analysis or fixed for morphological analysis.

### Histological analysis and imaging

Hematoxylin and eosin (H&E) staining of paraffin-embedded liver sections was performed as previously described (Gromovsky et al., 2017; Warrier et al., 2015; Brown et al., 2010; Brown et al., 2008a; Brown et al. 2008b). Histopathologic evaluation was scored in a blinded fashion by a board-certified pathologist with expertise in gastrointestinal/liver pathology (Daniela S. Allende – Cleveland Clinic). H&E slides were scanned using a Leica Aperio AT2 Slide Scanner (Leica Microsystems, GmbH, Wetzlar, Germany) and images were processed using ImageScope (Aperio, Software Version 12.1)

### Quantification of adipose tissue immune cell populations by flow cytometry

After 16-weeks of HFD-feeding, gonadal WAT was excised, washed with 1X PBS, and immediately placed into RPMI with 1mg/mL Type II Collagenase (Sigma Aldrich, St. Louis, MO) for 30min at 37° C with gentle agitation. Digested clumps of tissue were pressed through a 100um strainer and washed with 1X PBS. Cells were centrifuged at 500g for 10min; supernatant was aspirated. Cell pellet containing the stromal vascular fraction was resuspended in ACK Lysing Buffer (Life Technologies, Grandstand, NY) for 5min at room temperature. Cells were washed with 1X PBS and centrifuged at 300g for 10min. For flow cytometry, cells were resuspended in freshly prepared FACS buffer (1x PBS, 3% FBS) and aliquoted into 96 well plates. Cells were centrifuged at 830 x g for 4 minutes, resuspended in 50ul FACS buffer containing 0.5 μg anti-mouse CD16/CD32 mAb 2.4G2 (Mouse BD Fc Block, BD Pharmingen, San Diego, California), and incubated for 15 minutes 4° C. After blocking, cells were stained with a fluorochrome-conjugated antibody panel (antibodies described in the key resource table) for 30 minutes at 4° C in the dark. Cells were washed and centrifuged at 830 x g for 4 minutes twice with FACS buffer. Stained cells are resuspended in 200ul of 1% paraformaldehyde and kept in the dark at 4° C overnight. Stained cells were centrifuged at 830 x g for 5 minutes. Stained cells were resuspended in 300ul of FACS buffer, and data was collected on a Cytek Aurora full sprectrum (365-829 nm range) cytometer using SpectroFlo^®^ software(Cytek^®^ Biosciences, Fremont, California). Data collected on the Aurora were analyzed using FlowJo software (Tree Star, Inc., Ashland Oregon). Gating strategies are shown in Figure 4 – figure supplement 1.

### Immunoblotting

Whole tissue homogenates were made from tissues in a modified RIPA buffer as previously described (Schugar et al., 2017; Gromovsky et al., 2017; Warrier et al., 2015; Brown et al., 2010; Brown et al., 2008a; Brown et al. 2008b) that was supplemented with an additional 2% (w/v) sodium dodecylsulfate, and protein was quantified using the bicinchoninic (BCA) assay (Pierce). Proteins were separated by 4–12% SDS-PAGE, transferred to polyvinylidene difluoride (PVDF) membranes, and proteins were detected after incubation with specific antibodies as previously described (Schugar et al., 2017; Gromovsky et al., 2017; Warrier et al., 2015; Brown et al., 2010; Brown et al., 2008a; Brown et al. 2008b).

### RNA Isolation and bulk RNA sequencing

gWAT RNA was isolated using Trizol-Chlorofom extraction, gDNA removal by eliminator spin column (Qiagen), and isopropanol precipitation followed by ethanol clean-up. RNA quality was confirmed by Bioanalyzer (Agilent). RNA-SEQ libraries were generated using the Illumina mRNA TruSEQ Directional library kit and sequenced using an Illumina HiSEQ4000 (both according to the Manufacturer’s instructions). RNA sequencing was performed by the University of Chicago Genomics Facility. Raw sequencing data in the form of fastq files were transferred to and analyzed by the Bioinformatics Core at Case Western Reserve University. Fastq files were trimmed for quality and adapter sequences using TrimGalore! (version 0.6.5 Babraham Institute, https://github.com/FelixKrueger/TrimGalore) a wrapper script for CutAdapt and FastQC. Reads passing quality control were aligned to the mm10 mouse reference genome using STARAligner [cit] (version 2.5.3a) guided using the GENCODE gene annotation. Aligned reads were analyzed for differential gene expression using Cufflinks [cit] (version 2.2.1) which reports the fragments per kilobase of exon per million fragments mapped (FPKM) for each gene. Significant genes were identified using a significance cutoff of q-value <0.05 (FDR) and used as input for downstream analysis.

### Plasma and liver biochemistries

To determine the level of hepatic injury in mice fed HFD, plasma was used to analyze alanine aminotransferase (ALT) levels using enzymatic assays as previously described (Helsley et al., 2019). Extraction of liver lipids and quantification of total plasma and hepatic triglycerides, total cholesterol, and free cholesterol was conducted using enzymatic assays as described previously (Gromovsky et al., 2017; Warrier et al., 2015; Brown et al., 2010; Brown et al., 2008a; Brown et al. 2008b).

### Targeted quantification of phosphatidylinositol (PI) and lysophosphatidylinositol (LPI) lipids

Quantitation of LPI and PI species was performed as previously described (Helsley et al., 2019) Briefly, LPI and PI standards (LPI-16:0, LPI-18:0, LPI-18:1, LPI-20:4, PI-38:4) and the two internal standards (LPI-17:1-d31, PI-34:1-d31) were purchased Avanti Polar Lipids. HPLC grade water, methanol and acetonitrile were purchased from Thermos Fisher Scientific. Standard LPI and PI species at concentrations of 0, 5, 20, 100, 500 and 2000 ng/ml were prepared in 90% methanol containing 2 internal standards at the concentration of 500 ng/ml. The volume of 5 μl was injected into the Shimadzu LCMS-8050 for generating the internal standard calibration curves. A triple quadrupole mass spectrometer (Quantiva, Thermos Fisher Scientific, Waltham, MA, USA) was used for analysis of LPI and PI species. The mass spectrometer was coupled to the outlet of an UHPLC system (Vanquish, Thermos Fisher Scientific, Waltham, MA, USA), including an auto sampler with refrigerated sample compartment and inline vacuum degasser. The HPLC eluent was directly injected into the triple quadrupole mass spectrometer and the analytes were ionized at ESI negative mode. Analytes were quantified using Selected Reaction Monitoring (SRM) and the SRM transitions (m/z) were 571 → 255 for LPI-16:0, 599 → 283 for LPI-18:0, 597 → 281 for LPI-18:1, 619 → 303 for LPI-20:4, 885 → 241 for PI-38:4, 583 → 267 for internal standard LPI-17:1, and 866 → 281 for internal standard PI-34:1-d31. Xcalibur software was used to get the peak area for both the internal standards and LPI and PI species. The internal standard calibration curves were used to calculate the concentration of LPI and PI species in the samples.

### Untargeted lipidomics and oxylipin quantification

Lipidomics and oxylipin data were acquired at the UC Davis West Coast Metabolomics Center. For complex lipid analyses, reverse phased liquid chromatography-tandem mass spectrometry (RPLC-MS/MS) was used to perform lipidomic analysis and began by adding 100 μL run solvent (9 : 1 methanol/toluene (v/v)) to microcentrifuge tubes from the dried upper layer of extraction. Chromatography was performed using an Agilent 1290 UHPLC and mass spectra were collected with an Agilent 6550 QTOF mass spectrometer. An Acquity UPLC CSH C18 (100 mm × 2.1 mm, 1.7 μm particle size) column (Waters, Milford, MA) with an Acquity UPLC CSH C18 (5 mm × 1.2 mm, 1.7 μm particle size) pre-column (Waters, Milford, MA) was used with mobile phase A (6 : 4 acetonitrile/water (v/v)) and mobile phase B (9 : 1 isopropanol acetonitrile (v/v)). Mobile phases A and B were modified with 10 mM ammonium formate and 0.1% formic acid for positive mode ionization and 10 mM ammonium acetate for negative mode ionization. The LC gradient started at 15% B, increased to 30% B from 0–2 minutes, increased to 48% B from 2–2.5 minutes, increased to 82% B from 2.5–11 minutes, increased to 99% B from 11–11.5 minutes, held at 99% B from 11.5–12 minutes, returned to 15% B from 12–12.1 minutes and held at 15% B from 12.1–15 minutes. The autosampler was held at 4 °C and needle wash was performed before and after sample injections for 10 seconds with isopropanol. Injection volumes were 4 μL for both positive and negative mode ionization analyses. Spectral data was collected with a scan range of 120–1700 m/z. MS/MS fragmentation used data-dependent-acquisition (DDA) and was collected for the top 4 most abundant ions from each MS scan. LC-MS/MS data were processed using open source software MS-DIAL (Folz et al., 2021) (version 4.24) which performed peak picking, deisotoping, automated peak annotation, alignment and gap filling. Blank subtraction was performed by removing features that had a maximum sample intensity/average blank intensity ratio of less than 5. Adduct and duplicate features were flagged using Mass Spectral Feature List Optimizer (MS-FLO) (Folz et al., 2021). Data from each of the four analytical platforms (RPLC-MS/MS ESI+/−, HILIC-MS/MS ESI+/−) were processed separately and combined after data curation. No data normalization was performed because no trend in data intensities was observed from the internal standards during data acquisition. Peak height was used for all quantitation. Metabolite annotations were made using defined confidence levels (Folz et al., 2021) based on accurate mass, MS/MS library matching to experimental data, and retention time from authentic standards run on the same instrument. Tandem MS/MS libraries of the MassBank of North America (<u>MassBank.us</u>) and NIST17 (NIST, Gaithersburg, MD) were used for spectral matching. A manual curation of datasets was performed to reduce in-source fragment annotations identified by very similar RT and high correlation between features. Manual review of MS/MS matches was performed to remove poor spectral matches since false positive annotations can occur when automatically matching MS/MS from complex biological samples to large MS/MS spectral libraries (Folz et al., 2021). For oxylipin analyses, samples were added to a 96-well plate followed sequentially by 25 μL anti-oxidant solution, 25 μL of surrogate standards in methanol, 25 μL of CUDA and PHAU standards in methanol, and 125 μL acetonitrile/methanol 1:1. Plates were then vortexed for 30 s, centrifuged at 6 °C for 5 min at 15,000 rcf and filtered through a PVDC 0.2-micron filter plate. Plates were then sealed and kept at −20 °C until analysis. Extracted oxylipins were separated and quantified using a Waters i-Class Acquity UHPLC system coupled to a Sciex 6500+ QTRAP mass spectrometer operated in negative ionization mode. Oxylipins were quantified by targeted, retention time-specific, multiple reaction monitoring ion transitions. A total of 78 oxylipins were targeted, and targets that appeared in any sample above the signal-to-noise ratio of 3:1 were quantified. This resulted in quantified values for 27 n-6-related oxylipins, including 17 ARA-related and 10 LA-related oxylipins. There were 12 n-3- related oxylipins measured, which included 6 ALA-related, 4 DHA-related, and 2 EPA-related oxylipins. Oxylipins measured in a sample below the limit of quantitation (LOQ), defined as signal-to-noise ratio below 3:1, were converted to 10% of the LOQ.

### Statistical analyses

For the data shown in Figure 1 (hybrid mouse diversity panel) correlations and associated p values were calculated with the biweight midcorrelation, which is robust to outliers and associated p-value. Single comparisons between two groups were performed using two-tailed Student’s t tests with 95% confidence intervals. Comparisons involving multiple time points were assessed using an ordinary 1-way ANOVA followed by Tukey’s multiple comparisons test, or using a 2-way ANOVA followed by Sidak’s multiple comparisons test. All data presented as mean ± SEM. Values were considered significant at p < 0.05. *p< 0.05, **p<0.01, ***p<0.001 and ****p<0.0001. For all other figure panels each experiment consisted of a minimum of two biological replicates and data are presented as mean ± SD except where noted in the figure legend. All data were analyzed using either one-, two-, or three-way analysis of variance (ANOVA) where appropriate, followed by either Tukey’s or Student’s t-tests for post hoc analysis. Differences were considered significant at p <0.05. GraphPad Prism 9.2.0 (La Jolla, CA) was used for data analysis.

## ACKNOWLEDGEMENTS

This work was supported in part by National Institutes of Health grants R01 DK120679 (J.M.B.), P01 HL147823 (J.M.B.), P50 AA024333 (J.M.B.), U01 AA026938 (J.M.B.), R01 DK130227 (J.M.B.), R01 DK117850 (A.J.L.), and R01 HL148577 (A.J.L.). The authors would like to thank Dr. Philip Hawkins (Babraham Institute) for providing *Mboat7* knockout first mice. Some of the lipidomics and oxylipin data were acquired at the UC Davis West Coast Metabolomics Center.

## AUTHOR CONTRIBUTIONS

W.M. and J.M.B. planned the project, designed experiments, and analyzed data. W.M. and J.M.B. wrote the original draft of the manuscript; D.S.A. and A.J.L. helped design experiments and provided useful discussion directing collaborative aspects of the project; W.M., V.V., R.B., A.L.B., A.J.H., R.C.H., E.R.C., C.P., R.Z., D.S.A., A.J.L., and J.M.B. either conducted mouse experiments, performed biochemical workup of mouse tissues or analyzed data and aided in manuscript preparation. All authors were involved in the editing of the final manuscript.

## DATA AVAILABILITY

RNA sequencing data is in process at GEO Profiles at the National Center for Biotechnology Information (NCBI), and full accession will be available upon acceptance of this work. All materials, methods, and datasets included in this manuscript are readily available upon request.

## Abbreviations used

AA: arachidonic acid
ASO: antisense oligonucleotide
GWAS: genome wide association studies
LPI: lysophosphatidylinositol
LPIAT1: lysophosphatidylinositol acyltransferase 1
*MBOAT7*: membrane-bound O-acyltransferase 7
*Mboat7*^ASKO^: *Mboat7* adipocyte-specific knockout mice
*Mboat7*^HSKO^: *Mboat7* hepatocyte-specific knockout mice
MPO: morpholino oligonucleotides
NAFLD: non-alcoholic fatty liver disease
PC: phosphatidylcholine
PI: phosphatidylinositol
PIS: phosphatidylinositol synthase
SREBP-1c: sterol regulatory element-binding protein 1c
TAG: triacylglycerol.

## FIGURE LEGENDS

**Figure 2 – figure supplement 1.**
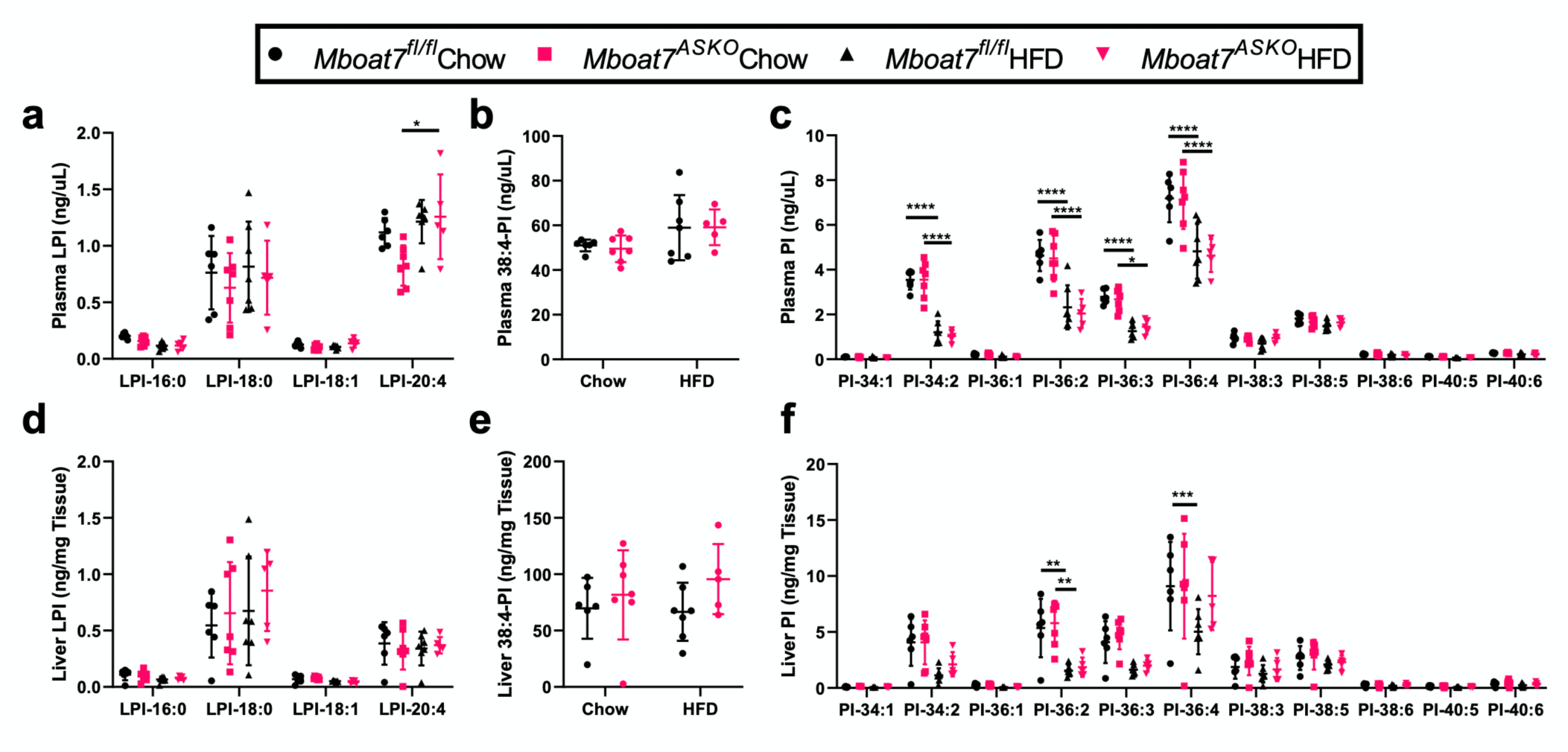
Plasma and Liver LPI and PI Levels are Not Altered in *Mboat7^ASKO^* Mice. Male control (*Mboat7^fl/fl^*) or adipocyte-specific Mboat7 knockout mice (*Mboat7^ASKO^*) were fed chow or high fat diet (HFD) for 20-weeks. Plasma LPI (**a**), PI-38:4 (**b**), and other PI species (**c**) were, were quantified via LC-MS in *Mboat7^fl/fl^* or *Mboat7^ASKO^* mice (n=5-7; \*\*\*\**P*≤0.0001; Two-way (**b**) or Three-way (**a,c**) ANOVA with Tukey’s *post-hoc* test). Liver LPI (**d**), PI-38:4 (**e**), and other PI species (**f**) were, were quantified via LC-MS in *Mboat7^fl/fl^* or *Mboat7^ASKO^*mice were fed chow or HFD for 20-weeks (n=5-7; \*\*\*\**P*≤0.0001; Two-way (**e**) or Three-way (**d,f**) ANOVA with Tukey’s *post-hoc* test). All data are presented as mean ± S.D.

**Figure 2 – figure supplement 2.**
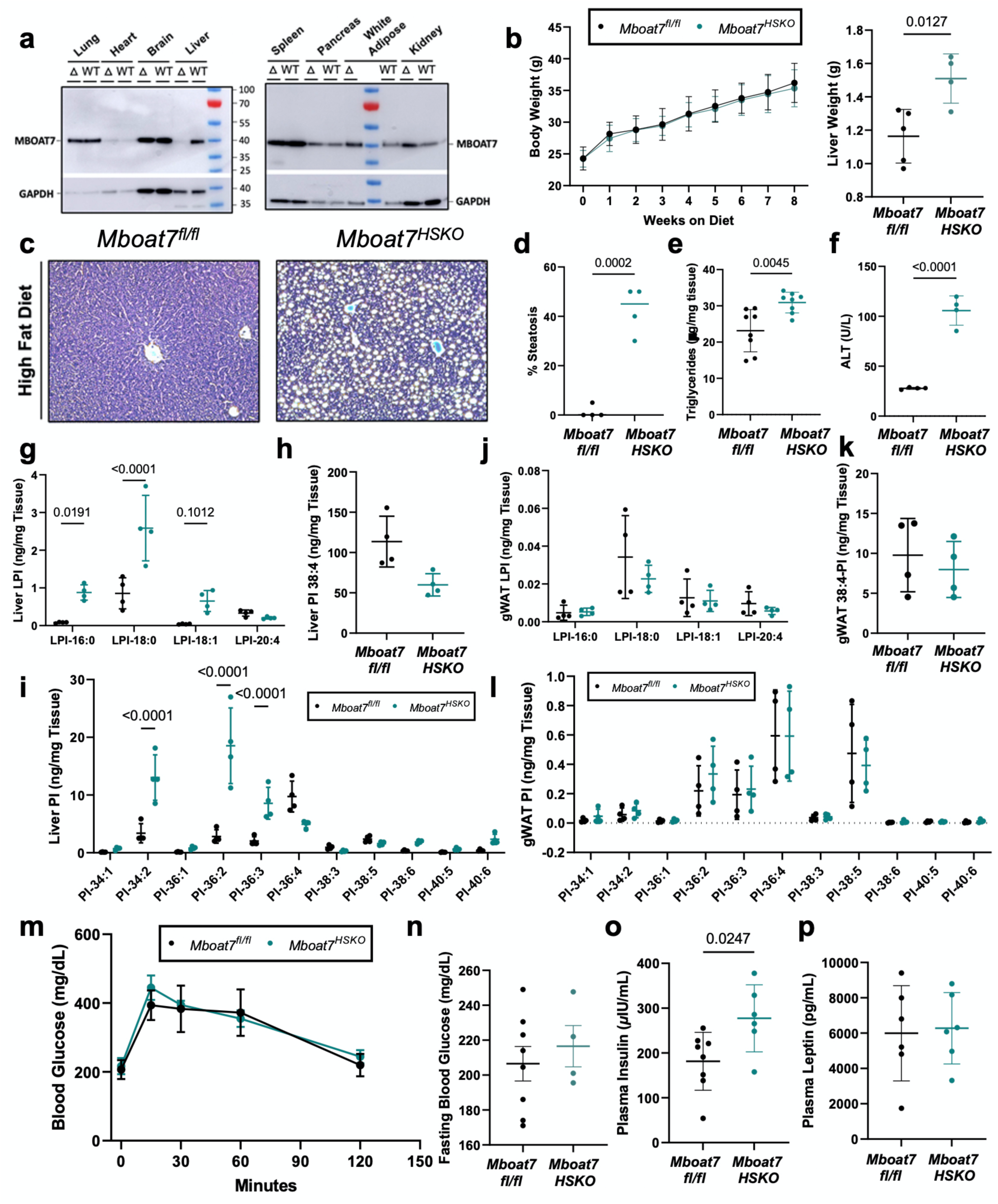
High Fat Diet Feeding in Hepatocyte-Specific *Mboat7* Knockout Mice (*Mboat7^HSKO^*) Results in Profound Fatty Liver Without Altering Glucose Tolerance. Male control (*Mboat7^fl/fl^*) or hepatocyte-specific Mboat7 knockout mice (*Mboat7^HSKO^*) were fed chow or high fat diet (HFD) for 8-weeks and metabolically phenotyped. (**a**) Western blots on microsomal fractions from hepatocyte-specific *Mboat7* deletion (*Mboat7^HSKO^*) tissues were probed for MBOAT7 and GAPDH to show a specific reduction of MBOAT7 protein in the liver. (**b-l**) *Mboat7^fl/fl^*or *Mboat7^HSKO^* fed an HFD for 10 weeks. (**b**) Body weight curve (n=6-8/group; Two-way ANOVA with Tukey’s *post-hoc* test). (**c**) Representative liver hematoxylin and eosin stained sections. 20x magnification. (**d**) Percent steatosis was quantified by a blinded pathologist (n=4/group; Two-sided Student’s t-test). Hepatic triglycerides (**e**) and plasma alanine aminotransferase (ALT) (**f**) were measured enzymatically (n=8) (**e**) or 4 (**f**); Two-sided Student’s t-test). Liver LPI (**g**), PI-38:4 (**h**), and other PI species (**i**) were, were quantified via LC-MS in *Mboat7^fl/fl^* or *Mboat7^HSKO^*mice were fed HFD for 10-weeks (n=4/group; Two-sided Student’s t-test (**h**) or Two-way ANOVA with Tukey’s *post-hoc* test (**g,i**)). gWAT LPI (**j**), PI-38:4 (**k**), and other PI species (**l**) were, were quantified via LC-MS in *Mboat7^fl/fl^* or *Mboat7^HSKO^* mice were fed chow or HFD for 10-weeks (n=5-7; \*\*\*\**P*≤0.0001; Two-sided Student’s t-Test (**k**) or Two-way ANOVA with Tukey’s *post-hoc* test (**j,l**). (**m-n**) *Mboat7^fl/fl^* or *Mboat7^HSKO^*mice were fed an HFD for 3-4 weeks and then underwent an intraperitoneal glucose tolerance test (GTT). (**m**) Plasma glucose levels were measured (in duplicate or triplicate at each time point) throughout the GTT (n=4-8; Two-way ANOVA with Tukey’s *post-hoc* test). (**n**) Fasting blood glucose was measured after a four hour fast (time=0 minutes for GTT) (n=4-8; Two-sided Student’s t-Test). Fasting plasma insulin (**o**) and leptin (**p**) was measured in *Mboat7^fl/fl^* or *Mboat7^HSKO^* fed an HFD for 10 weeks. Data in (**a-l,o,p**) are presented as mean ± S.D. Data in (**m,n**) are presented as mean ± S.E. M. (blood glucose readings during GTT were taken in duplicate or triplicate if duplicate measures varied by >10%).

**Figure 2 – figure supplement 3.**
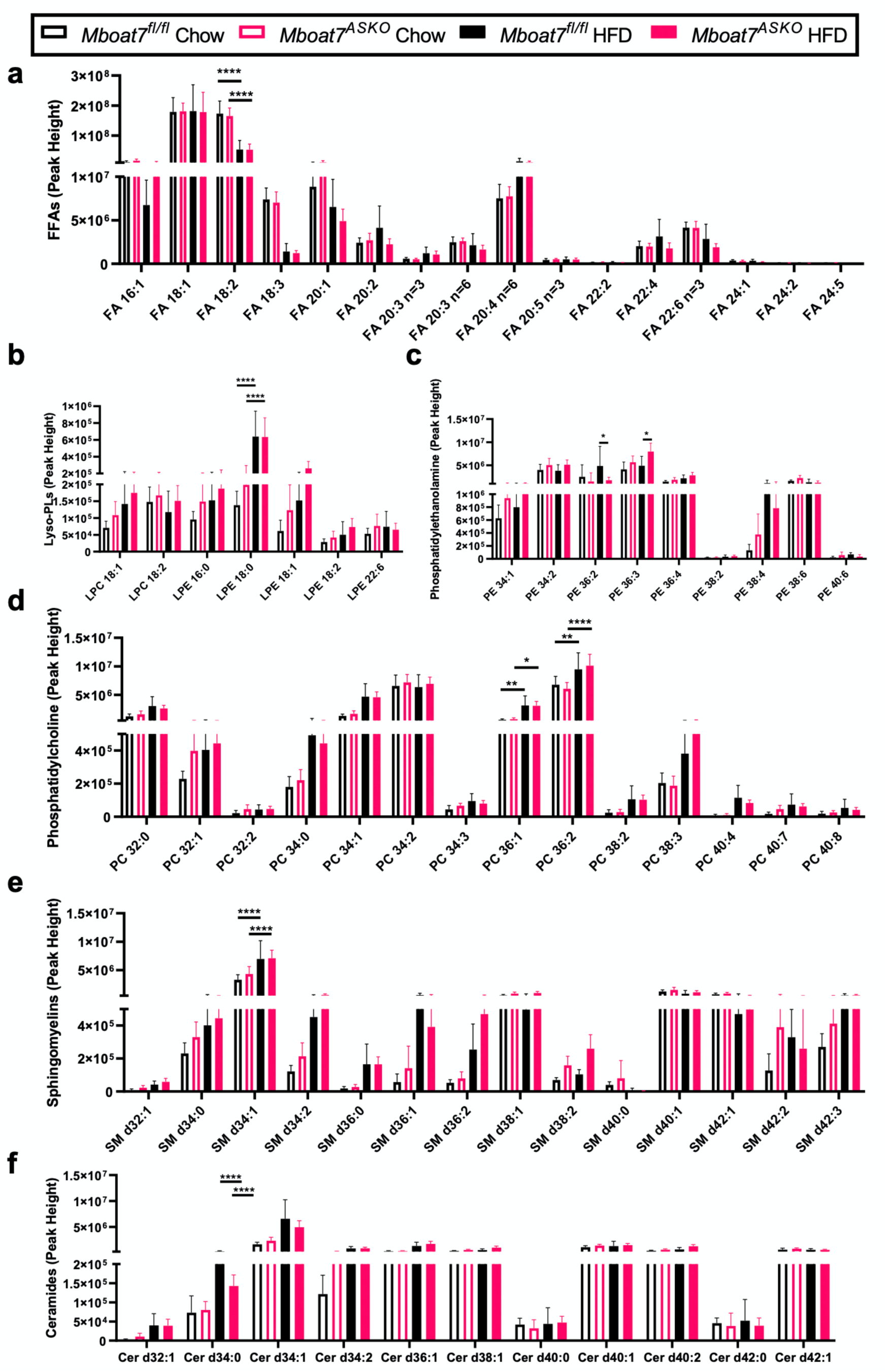
Adipose Tissue Free Fatty Acids, Glycerophospholipids, Sphingomyelins, and Ceramides are Not Altered in *Mboat7^ASKO^* mice. Male control (*Mboat7^fl/fl^*) or adipocyte-specific Mboat7 knockout mice (*Mboat7^ASKO^*) were fed chow or high fat diet (HFD) for 20-weeks. Gonadal adipose tissue (gWAT) free fatty acids (FFAs) (**a**), Lysophospholipids (LPLs) (**b**), Phosphatidyl-ethanolamines (PEs) (**c**), Phosphatidylcholines (PCs) (**d**), Sphingomyelins (SMs) (**e**), and Ceramides (Cers) (**f**) were quantified in *Mboat7^fl/fl^* or *Mboat7^ASKO^* mice were fed chow or HFD for 20-weeks via liquid chromatography mass spectrometry (n=5/group; Three-way ANOVA with Tukey’s *post-hoc* test). All data are presented as mean ± S.D.

**Figure 2 – figure supplement 4.**
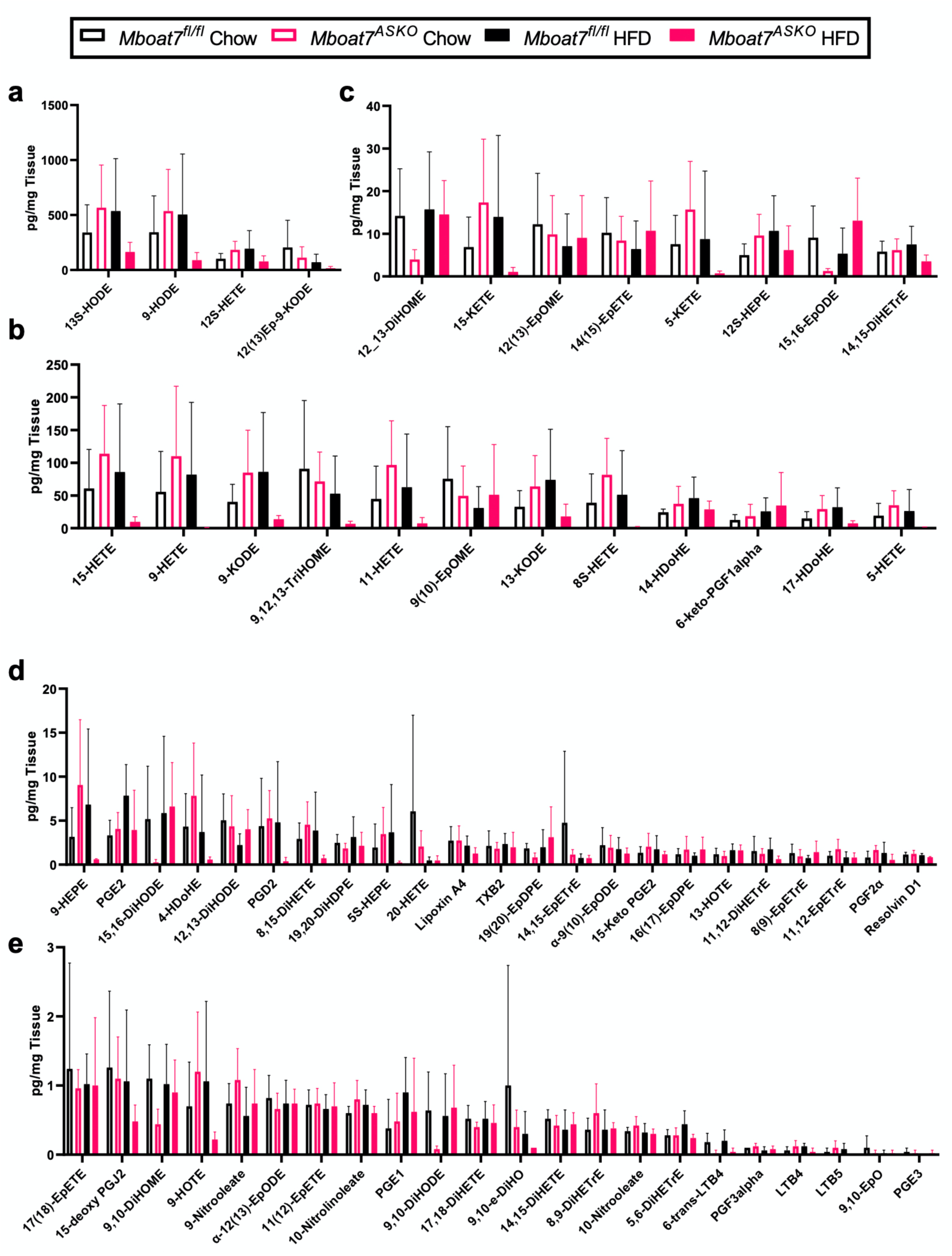
Gonadal White Adipose tissue (gWAT) Oxylipins are Not Altered in *Mboat7^ASKO^* mice. (**a-e**) Male control (*Mboat7^fl/fl^*) or adipocyte-specific Mboat7 knockout mice (*Mboat7^ASKO^*) were fed chow or high fat diet (HFD) for 20-weeks. Molecular species of oxylipins were quantified in gWAT of *Mboat7^fl/fl^* or *Mboat7^ASKO^* mice were fed chow or HFD for 20-weeks by liquid chromatography mass spectrometry (n=5/group); Three-way ANOVA with Tukey’s *post-hoc* test). All data are presented as mean ± S.D.

**Figure 2 – figure supplement 5.**
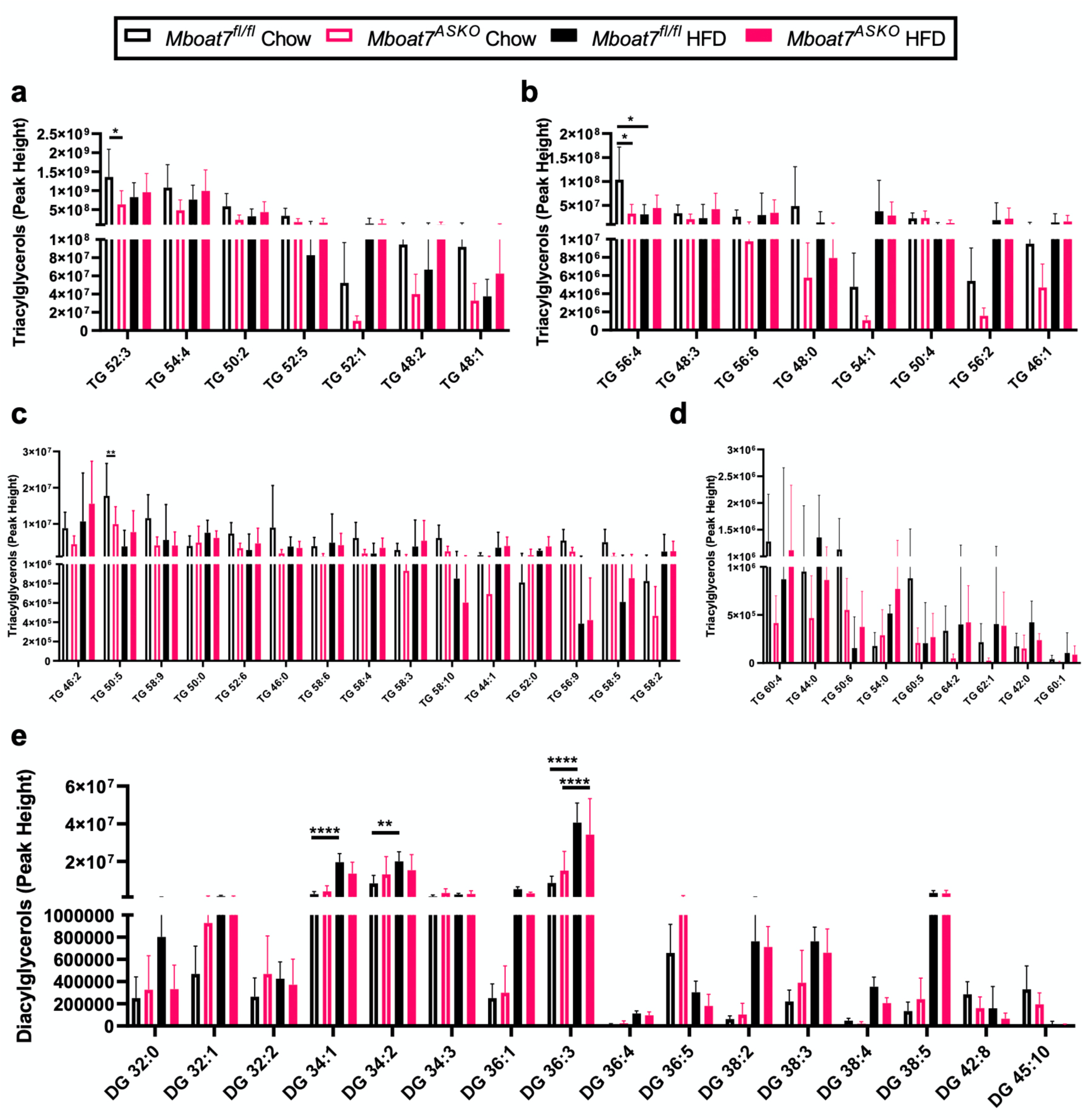
Adipose tissue Diacylglycerol (DAG) and Triacylglycerol (TAG) are Not Altered in *Mboat7^ASKO^* Mice. Male control (*Mboat7^fl/fl^*) or adipocyte-specific Mboat7 knockout mice (*Mboat7^ASKO^*) were fed chow or high fat diet (HFD) for 20-weeks. (**a-d**) Molecular species of triacylglycerols (TAG) were quantified via liquid chromatography mass spectrometry (n=5/group; Three-way ANOVA with Tukey’s *post-hoc* test). (**e**) Molecular species of diacylglycerols (DAG) were quantified in gWAT of *Mboat7^fl/fl^* or *Mboat7^ASKO^* mice were fed chow or HFD for 20-weeks via liquid chromatography mass spectrometry (n=5/group; Three-way ANOVA with Tukey’s *post-hoc* test). All data are presented as mean ± S.D.

**Figure 2 – figure supplement 6.**
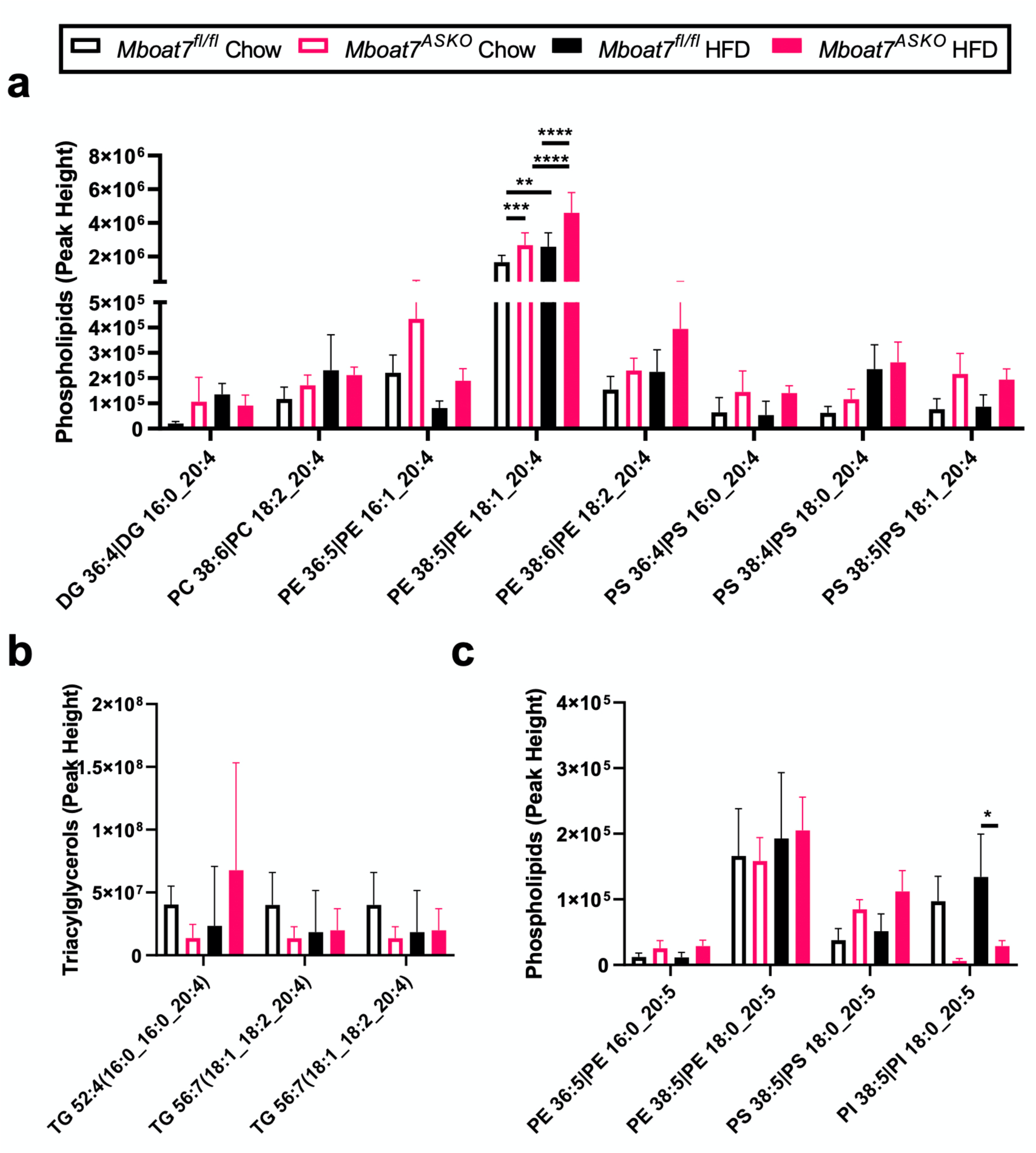
Adipocyte-Specific *Mboat7* Deletion Alters Select Arachidonic Acid and Eicosapentaenoic Acid Containing Lipids in Adipose Tissue. Male control (*Mboat7^fl/fl^*) or adipocyte-specific Mboat7 knockout mice (*Mboat7^ASKO^*) were fed chow or high fat diet (HFD) for 20-weeks. Phospholipids **(a,c)**, diacylglycerols (DAG) **(a)**, and triacylglycerols (TG) **(b)** containing arachidonic acid (20:4) were quantified in gWAT of *Mboat7^fl/fl^*or *Mboat7^ASKO^* mice were quantified by liquid chromatography mass spectrometry (n=5/group; Three-way ANOVA with Tukey’s *post-hoc* test). (**c**) Phospholipids containing eicosapentaenoic acid (20:5) were quantified in gWAT of *Mboat7^fl/fl^* or *Mboat7^ASKO^* mice were fed chow or HFD for 20-weeks by the West Coast Metabolomics’ complex lipid panel (n=5/group; Three-way ANOVA with Tukey’s *post-hoc* test). All data are presented as mean ± S.D.

**Figure 3 – figure supplement 1.**
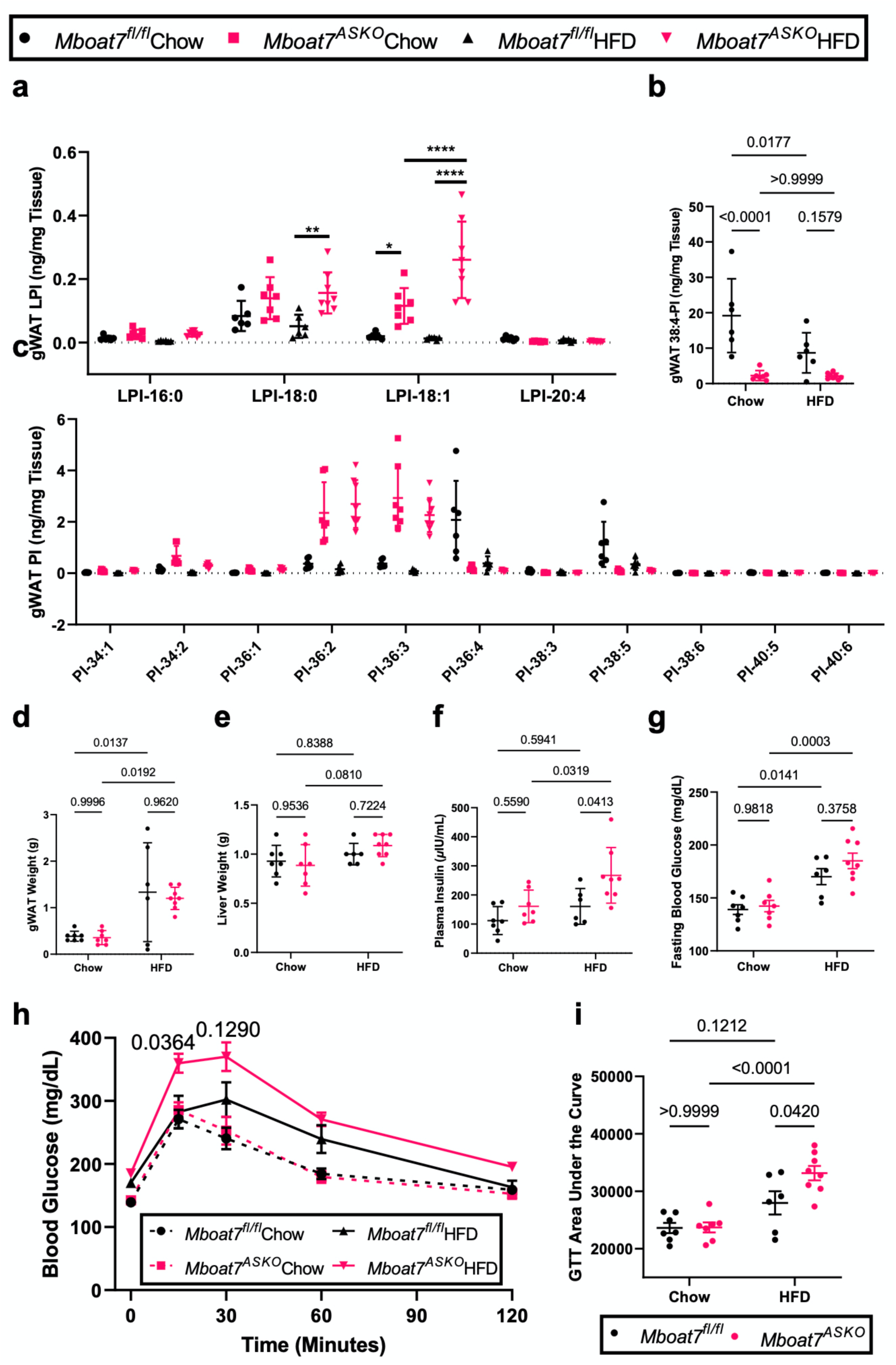
Female *Mboat7^ASKO^*Display Altered LPI/PI Balance in Adipose Tissue and Have Impaired Glucose Tolerance. Gonadal white adipose tissue (gWAT) Lysophosphatidylinositol (LPI) (**a**) and phosphatidylinositol (PI) species, including the MBOAT7 product PI-38:4 (**b**) and others (**c**), were quantified via LC-MS in *Mboat7^fl/fl^* or *Mboat7^ASKO^*mice were fed chow or HFD for 20-weeks (n=5-7; \*\*\*\**P*≤0.0001; Two-way (**b**) or Three-way (**a,c**) ANOVA with Tukey’s *post-hoc* test). gWAT (**d**) and liver (**e**) weight measurements from female *Mboat7^fl/fl^*or *Mboat7^ASKO^* mice fed Chow and HFD for 20 weeks (n=6-8; Two-way ANOVA with Tukey’s *post-hoc* test). (**f**) Fasting plasma insulin was measured in *Mboat7^fl/fl^*or *Mboat7^ASKO^* mice that were fed a chow or HFD for 20 weeks (n=6-8; Two-way ANOVA with Tukey’s *post-hoc* test). (**g-i**) Female *Mboat7^fl/fl^* or *Mboat7^ASKO^* mice were fed a chow or HFD for 12 weeks and then underwent an intraperitoneal glucose tolerance test (GTT). (**g**) Fasting blood glucose was measured after a four hour fast (time=0 minutes for GTT) (n=6-8; Two-way ANOVA with Tukey’s *post-hoc* test). (**h**) Plasma glucose levels were measured (in duplicate or triplicate at each timepoint) throughout the GTT (n=6-8; Three-way ANOVA with Tukey’s *post-hoc* test). (**i**) Area under the curve was calculated for each mouse throughout the GTT (n=6-8; Two-way ANOVA with Tukey’s *post-hoc* test). Data in (**a-f**) are presented as mean ± S.D. Data in (**g-i**) are presented as mean ± S.E. M. (blood glucose readings during GTT were taken in duplicate or triplicate if duplicate measures varied by >10%).

**Figure 3 – figure supplement 2.**
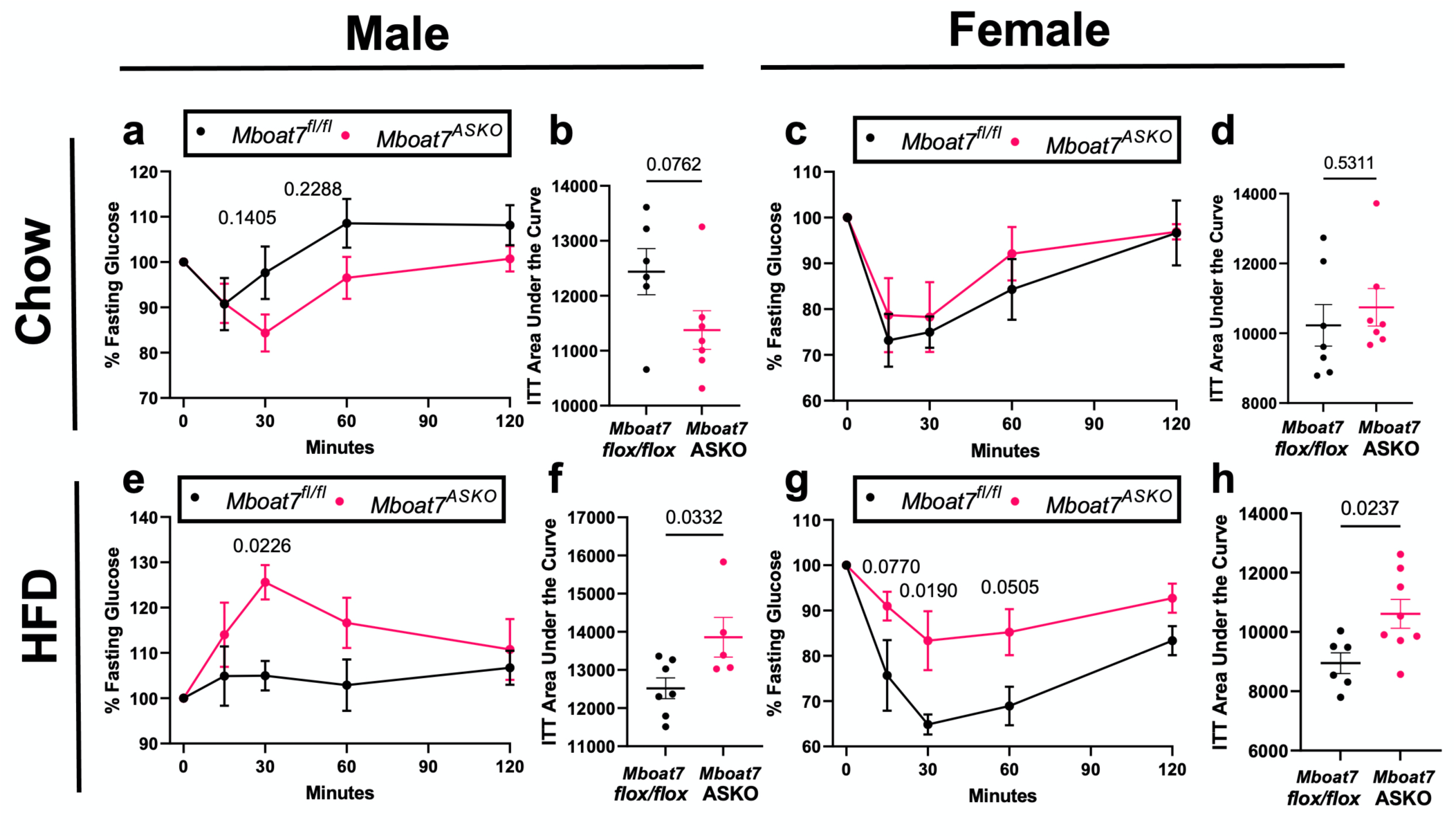
Adipocyte-Specific *Mboat7* Deletion (*Mboat7^ASKO^*) Promotes Insulin Resistance. (**a-c**) Male or female control (*Mboat7^fl/fl^*) or adipocyte-specific Mboat7 knockout mice (*Mboat7^ASKO^*) were fed a chow or HFD for 14 weeks and then underwent an insulin tolerance test (ITT). Plasma glucose levels were measured (in duplicate or triplicate at each timepoint) throughout the ITT and area under the curve (AUC) was calculated (n=5-7; Three-way ANOVA with Tukey’s *post-hoc* test).

**Figure 3 – figure supplement 3.**
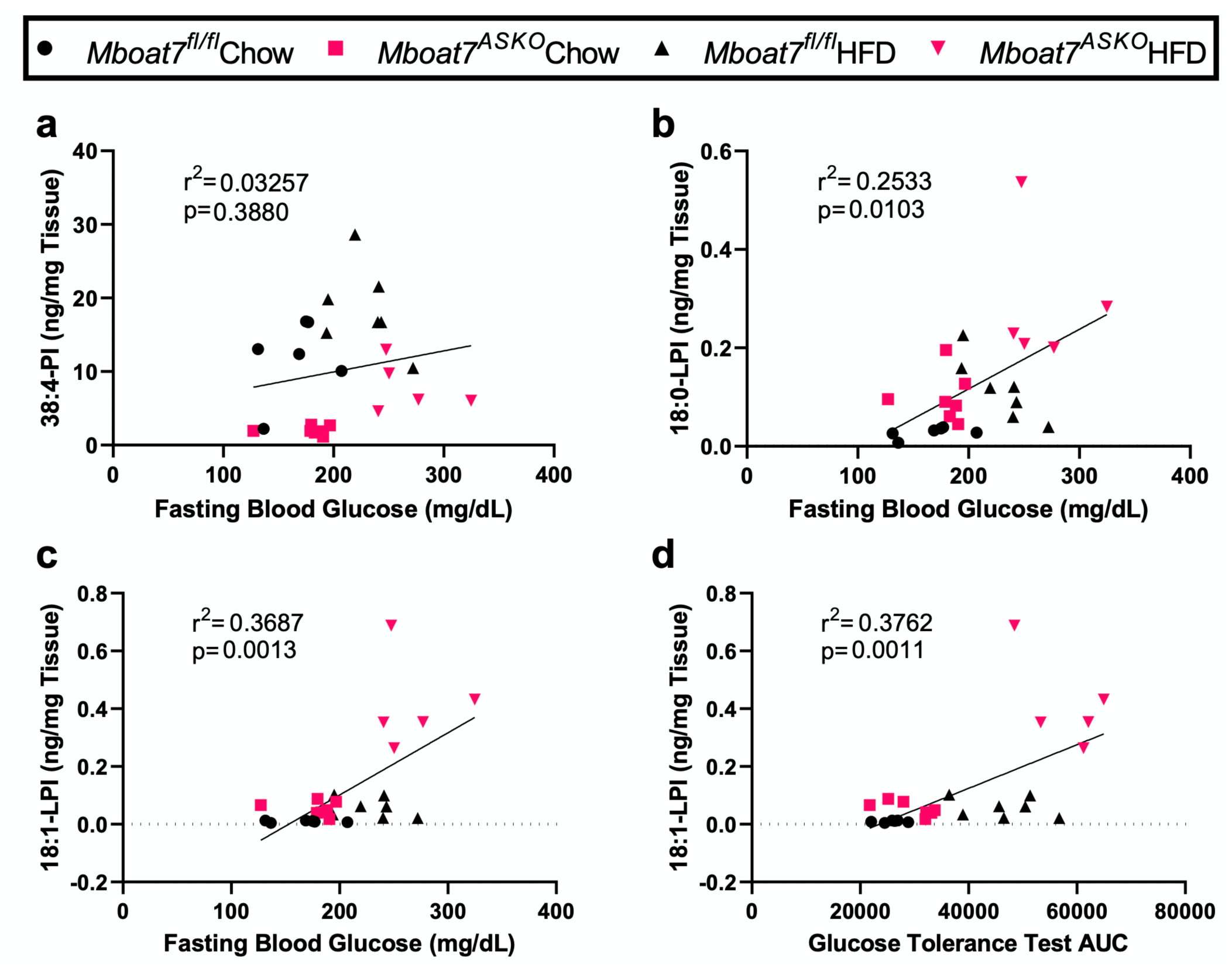
Adipose Tissue LPI Levels Correlate with Systemic Glucose Tolerance. Male control (*Mboat7^fl/fl^*) or adipocyte-specific Mboat7 knockout mice (*Mboat7^ASKO^*) were fed chow or high fat diet (HFD) for 20-weeks and correlation analysis was performed for glucose phenotypes and MBOAT7 substrate lysophosphatidylinositols (LPI) and product phosphatidylinositol (PI) lipids. (**a**) gWAT PI-38:4 at end of the experiment vs Fasting Blood Glucose at GTT. gWAT LPI-18:0 (**b**) and LPI-18:1 (**c**) at end of the experiment vs Fasting Blood Glucose at GTT. (**d**) gWAT LPI-18:1 at end of the experiment vs GTT Area under the curve.

**Figure 4 – figure supplement 1.**
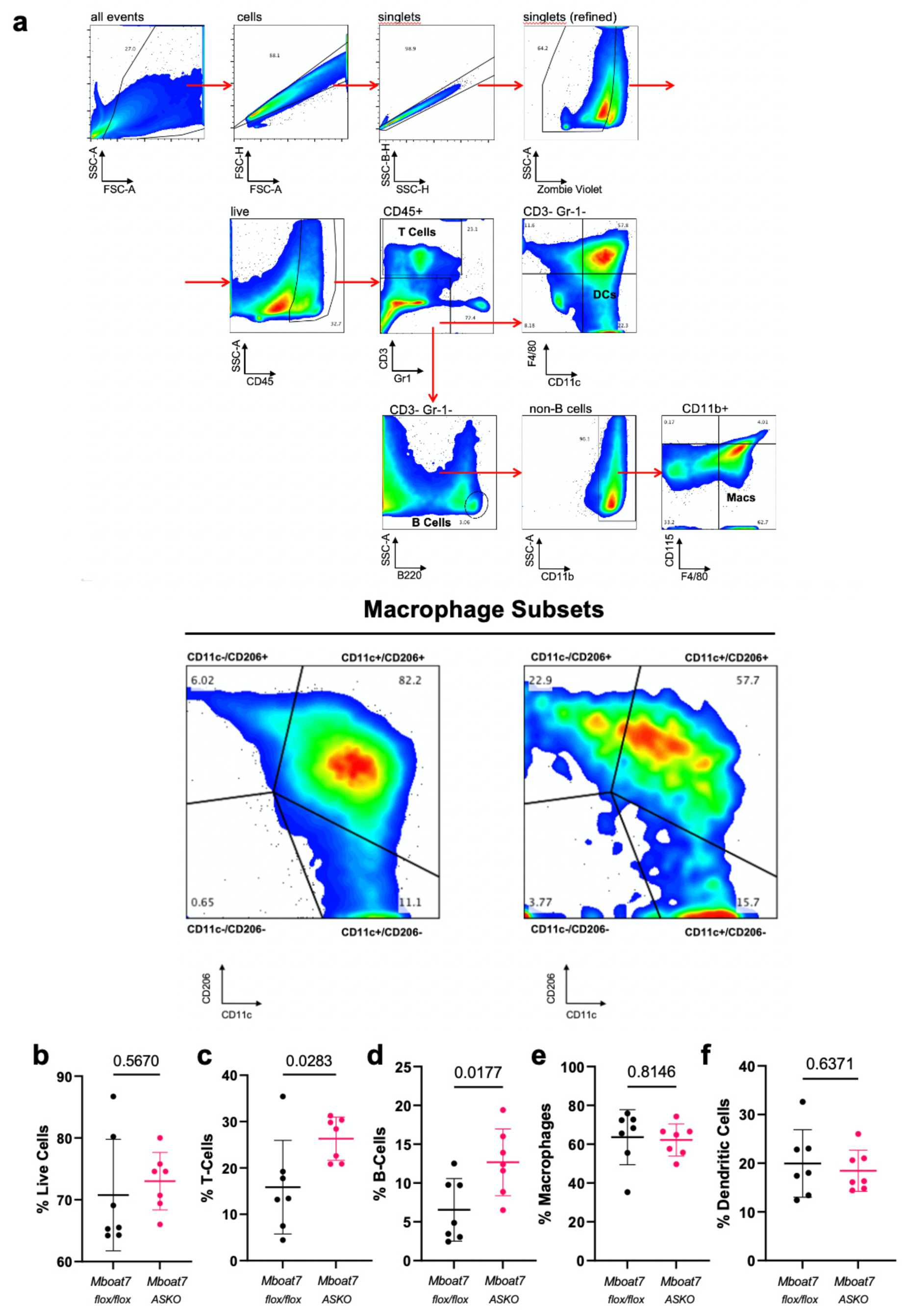
Adipocyte-Specific *Mboat7* Deletion (*Mboat7^ASKO^*) Reorganizes White Adipose Tissue Immune Cell Populations. Male control (*Mboat7^fl/fl^*) or adipocyte-specific Mboat7 knockout mice (*Mboat7^ASKO^*) were fed chow or high fat diet (HFD) for 20-weeks. The stromal vascular fraction of gonadal white adipose tissue (gWAT) was subjected to flow cytometric analysis of immune cell populations. **(a,b)** Gating strategy used for identifying subsets of adipose tissue macrophage populations. The percentage of live cells **(b)**, T-cells **(c)**, B-cells **(d)**, macrophages **(e)**, or dendritic cells **(f)** were quantified via flow cytometry.

